# Nucleolar detention of NONO shields DNA double-strand breaks from aberrant transcripts

**DOI:** 10.1101/2023.07.14.548979

**Authors:** Barbara Trifault, Victoria Mamontova, Giacomo Cossa, Sabina Ganskih, Yuanjie Wei, Julia Hofstetter, Pranjali Bhandare, Apoorva Baluapuri, Daniel Solvie, Carsten P. Ade, Peter Gallant, Elmar Wolf, Mathias Munschauer, Kaspar Burger

## Abstract

RNA-binding proteins (RBPs) stimulate the DNA damage response (DDR). The RBP NONO marks nuclear paraspeckles in unperturbed cells and undergoes poorly understood re-localisation to the nucleolus upon induction of DNA double-strand breaks (DSBs). Here we show that treatment with the topoisomerase-II inhibitor etoposide stimulates the production of RNA polymerase II-dependent, DNA damage-induced nucleolar antisense RNAs (diNARs) in human cells. diNARs originate from the nucleolar intergenic spacer and tether NONO to the nucleolus via its RRM1 domain. NONO occupancy at protein-coding gene promoters is reduced by etoposide, which attenuates pre-mRNA synthesis, enhances NONO binding to pre-mRNA transcripts and is accompanied by nucleolar detention of such transcripts. The depletion or mutation of NONO interferes with detention and prolongs DSB signaling. Together, we describe a nucleolar DDR pathway that shields NONO and aberrant transcripts from DSBs to promote DNA repair.

## Introduction

Genome stability requires the faithful inheritance of genetic information. The DNA damage response (DDR) recognizes and repairs DNA lesions to maintain genome stability(Jackson and Bartek 2009; Ciccia and Elledge 2010). Kinases like *Ataxia-telangiectasia mutated* (ATM) inhibit unscheduled RNA synthesis to suppress DNA double-strand breaks (DSBs) and promote DSB repair (DSBR) (Blackford and Jackson 2017; Machour and Ayoub 2020). However, transcripts and RNA-binding proteins (RBPs) emerge as regulators of the DDR an also stimulate DSBR (Duterte et al. 2014; Michelini et al. 2018; Burger et al. 2019; Zong et al. 2020). The *Drosophila* behavior/human splicing (DBHS) protein family member NONO associates with actively transcribed chromatin and paraspeckles in unperturbed cells, and participates in DSBR, for instance by stimulating non-homologous end joining (Shav-Tal and Zipori 2002; Krietsch et al. 2012; Knott et al. 2016; Jaafar et al. 2017; Wang et al. 2022). Interestingly, NONO accumulates in condensates induced by transcription inhibition to suppress gene fusions, but also undergoes poorly understood re-localisation to the nucleolus upon DNA damage (Moore et al. 2011; Yasuhara et al. 2022).

Here we show that the topoisomerase-II inhibitor etoposide stimulates a nucleolar DDR pathway, which involves RNA polymerase II (RNAPII)-dependent, DNA damage-induced nucleolar antisense RNAs (diNARs) that form nucleolar DNA-RNA-hybrids (R-loops) and deplete NONO from protein-coding gene promoters, which reduces pre-mRNA synthesis, detains aberrant transcripts and stimulates DSBR.

## Results

### The RRM1 domain facilitates NONO nucleolar re-localisation

We showed previously that treatment with etoposide enriches NONO in non-disintegrated nucleoli (Trifault et al. 2022) and confirmed this by costaining of DBHS proteins NONO, SFPQ or PSPC1 with nucleophosmin (NPM1) (Supplemental Fig. S1A,B). Etoposide treatment increased the ser-139 phosphorylated histone H2.X variant (γH2A.X) >5-fold, but not DBHS proteins (Supplemental Fig. S1C). Next, we tested if NONO re-localisation is induced by nucleolar DSBs. We employed the 4-hydroxytamoxifen (4-OHT)-inducible endonucleases I-PpoI, which cleaves 28S ribosomal (r)DNA (van Sluis and McStay 2019). We observed signals for γH2A.X and 53BP1, but not NONO around disintegrated nucleoli upon 4-OHT treatment (Supplemental Fig. S2A,B), suggesting that nucleoplasmic DSBs trigger NONO nucleolar re-localisation. NONO RNA recognition motifs 1/2 (RRM1/2) mediate binding to nucleic acids (Knott et al. 2016). To test, which domain confers nucleolar re-localisation, we created HA-NONO mutants (Supplemental Fig. S2C), monitored their expression (Supplemental Fig. S2D), and assessed their localisation (Fig. 1A, Supplemental Fig. S2E). Costaining of mutants with fibrillarin revealed that full length (FL) HA-NONO, the RRM1 deletion mutant (ΔRRM1) and the carboxy-terminal deletion mutant (ΔC-ter) localised in the nucleoplasm in unperturbed cells. Upon incubation with etoposide, FL and ΔC-ter displayed pan-nuclear localisation and co-staining with fibrillarin, which was impaired by ΔRRM1. Thus, NONO RRM1 confers nucleolar re-localisation.

**Figure 1.**
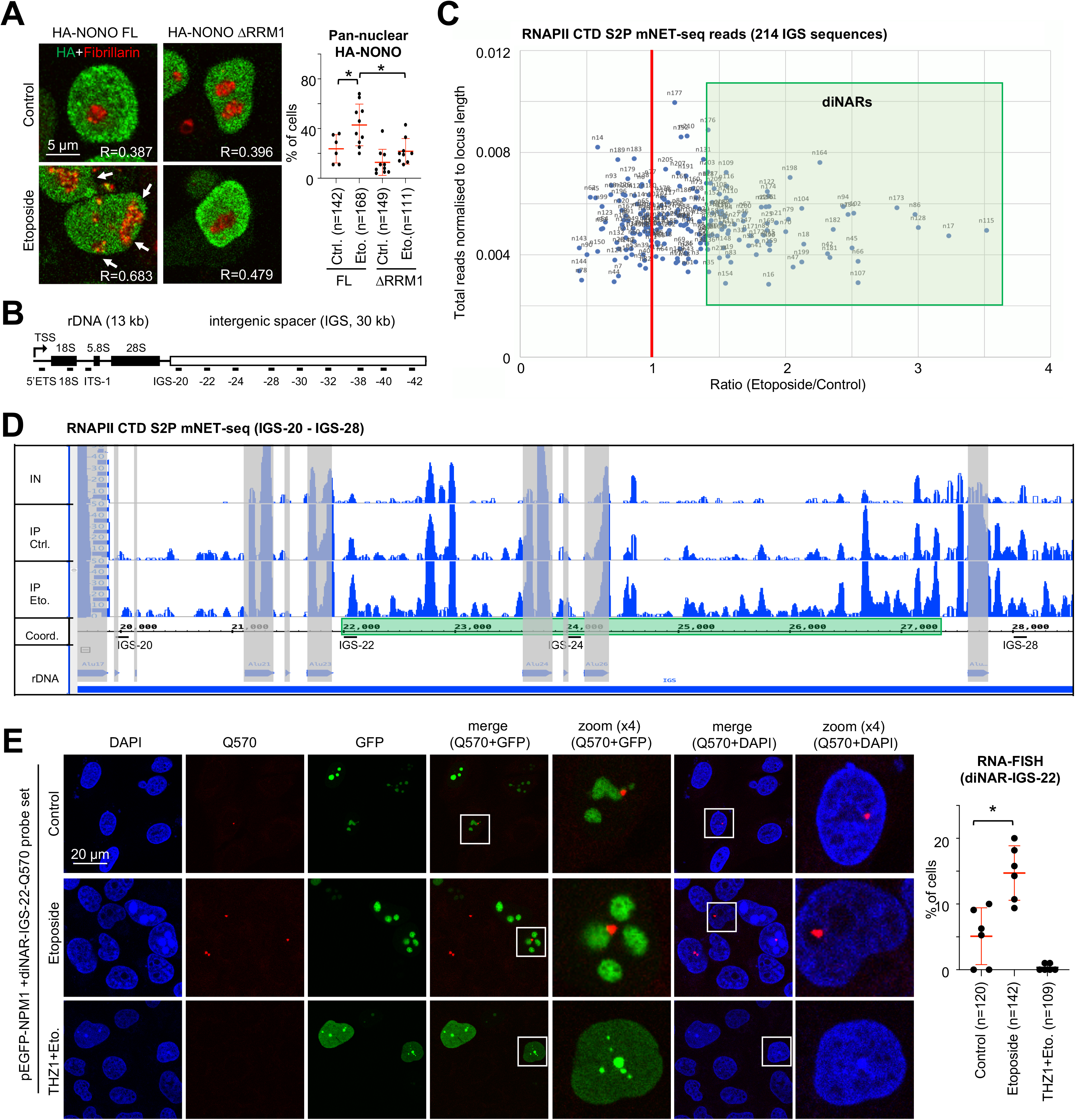
DNA damage induces RNAPII-dependent nucleolar transcripts in U2OS cells. (*A*) Imaging (left) and quantitation (right) of HA-NONO variants and fibrillarin. Arrowhead, colocalisation; R=Pearson correlation. n, number of cells. Each dot represents % of cells with pan-nuclear HA signals as average from one acquisition. (*B*) Scheme of human ribosomal (r)DNA array (∼80 repeats on chr. 13, 14, 15, 21, 22). Intergenic spacer (IGS) 20 to 42, probe positions in kb downstream of rDNA transcriptional start site (TSS). (*C*) Scatter plot displaying mNET-seq IGS reads. (*D*) mNET-seq browser tracks for IGS consensus region 20-28 from inputs (IN, merged) or after immunoprecipitation (IP) with CTD S2P-selective antibody ±etoposide. Grey, *Alu* element; green, induced region. (*E*) Imaging of GFP-NPM1 and Quasar570 RNA-FISH signals originating at IGS-22. White box, zoom (left) and quantitation (right). n, number of cells. Each dot represents % of cells with Quasar570-positive signals as average from two acquisitions. *, p-value <0.05; **, p-value <0.001; two-tailed t-test; n.d., not detected. Error bar, mean ±SD. Representative images are shown.

### RNAPII produces DNA damage-induced nucleolar transcripts at distinct IGS loci

Non-ribosomal, nucleolar transcripts maintain homeostasis by sequestration of RBPs (Mamontova et al. 2021; Feng and Manley 2022). Intriguingly, carboxy-terminal domain ser-2 phosphorylated RNA polymerase II (CTD S2P) synthesises antisense intergenic ncRNA (asincRNA) on nucleolar chromatin to regulate RNA polymerase I (RNAPI) in unperturbed cells (Abraham et al. 2020). We hypothesised that the DDR modulates nucleolar RNAPII activity and applied mammalian nascent elongation transcript sequencing (mNET-seq) to profile RNAPII-associated transcripts. First, we confirmed enrichment of RNAPII and associated transcripts upon immunoselection (Supplemental Fig. S3A). mNET-seq revealed that etoposide treatment elevated CTD S2P mNET-seq reads for about 25% of the 214 individually mapped intergenic spacer (IGS) sequences 2-3-fold, which we termed DNA damage-induced nucleolar antisense RNAs (diNARs) (Fig. 1B,C). Pair-wise comparison further suggested a locus-specific increase in diNARs at IGS loci 22, 30 and 38 (Supplemental Fig. S3B), which was also visualised on browser tracks (Fig. 1D, Supplemental Fig. S3C-E). Next, we used chromatin immunoprecipitation (ChIP) to assess CTD S2P occupancy. Treatment with etoposide, but not preincubation with the transcriptional kinase inhibitor THZ1, elevated CTD S2P signals at IGS loci 22, 30 and 38 (Supplemental Fig. S3F). Importantly, etoposide had little impact on CTD S2P marks (Supplemental Fig. S3G). Onset of antisense transcripts was validated by RT-qPCR with antisense-specific forward primers (Supplemental Fig. S3H). Next, we performed RNA-FISH to visualise prominently induced diNAR-IGS-22 and observed an increased number of cells comprising nucleolar RNA-FISH signals upon etoposide treatment (Fig. 1E, Supplemental Fig. S3I). We conclude that nucleolar CTD S2P produces diNARs.

### diNARs form nucleolar R-loops to promote NONO re-localisation

Nucleolar asincRNA form R-loops to shield RNAPI from the IGS (Abraham et al. 2020). As asincRNA-coding IGS loci and diNAR-encoding regions overlapp, we asked if nucleolar R-loops mediate NONO nucleolar re-localisation. We used S9.6 and NONO antibodies in DNA-RNA immunoprecipitation (DRIP) and NONO ChIP experiments. For S9.6 validation, we employed U2OS DIvA cells, which express the 4-OHT-inducible endonuclease AsiSI to induce DSBs (Clouaire et al. 2018). We assessed S9.6 reactivity at the R-loop-forming DS1 site (*RBMXL1* promoter) (Clouaire et al. 2018). We detected DRIP signals at DS1 in the presence of 4-OHT, which were sensitive to RNaseH digestion (Supplemental Fig. S4A). AsiSI cleavage was confirmed by imaging of γH2A.X and 53BP1 foci (Supplemental Fig. S4B). Next, we determined R-loop levels on nucleolar chromatin (Fig. 2A). We found that etoposide increased DRIP signals across the body of the IGS. To test if the formation of nucleolar R-loops correlates with elevated occupancy of NONO on nucleolar chromatin we performed NONO ChIP assays. For validation, we assessed NONO occupancy at DS1 in the absence or presence of NONO-targeting shRNA (Supplemental Fig. S4C). NONO occupancy at DS1 was sensitive to NONO depletion (Supplemental Fig. S4D). Next, we measured NONO occupancy on nucleolar chromatin. NONO ChIP signals were detectable on rDNA, but not responsive to etoposide (Supplemental Fig. S4E). On the IGS, however, NONO occupancy was modestly increased upon etoposide treatment, in particular after two hours of chase and at regions that displayed increased levels of diNARs and R-loops (Fig. 2B). To test if nucleolar R-loops promote NONO re-localisation, we overexpressed V5-tagged RNaseH1 and imaged NONO localisation (Fig. 2C). The etoposide-induced nucleolar re-localisation of NONO was impaired in cells that comparably express V5-RNaseH1 in the absence or presence of etoposide (Supplemental Fig. S4F). The etoposide-induced accumulation of NONO at IGS loci 22, 30, and 38 was also sensitive to overexpression of GFP-RNaseH1 (Fig. 2D). To asses if NONO binds R-loop-forming IGS loci directly, we performed pull-down assays with recombinant NONO (rec-NONO) and end-labeled DNA-RNA chimeras (gapmers). Gapmers were designed with sequence complementary to diNAR-encoding region IGS-22, or IGS-20 control, to mimic single-stranded DNA within R-loops. When incubating rec-NONO with immobilised biotinylated gapmers, we found that gapmer-22 enriched rec-NONO >2-fold stronger than gapmer-20 (Supplemental Fig. S4G). Next, we immobilised FL or ΔRRM1 HA-NONO variants on beads and incubated them with radio-labeled gapmers (Fig. 2E, Supplemental Fig. S4H). Immobilised FL, but not ΔRRM1, enriched gapmer-22 >2-fold stronger than gapmer-20. Thus, etoposide induces nucleolar R-loops to promote NONO nucleolar re-localisation.

**Figure 2.**
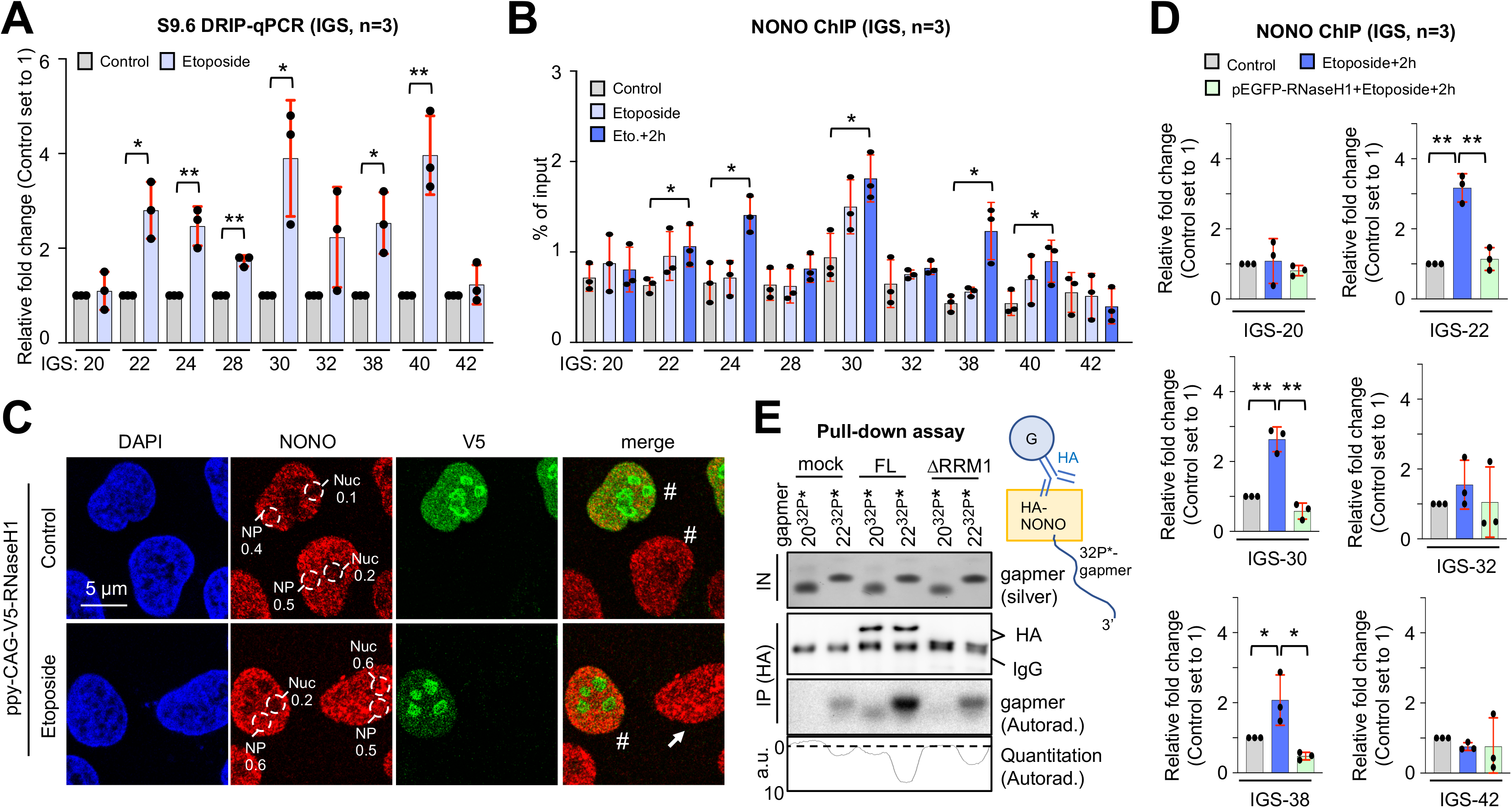
R-loop formation and NONO IGS occupancy correlate with diNAR synthesis and NONO nucleolar re-localisation in U2OS cells. (*A*) DRIP-qPCR using S9.6 antibody and region-specific primers. (*B*) NONO ChIP using site-specific primers. (*C*) Imaging of NONO and V5-RNaseH1. Arrowhead, pan-nuclear; #, nucleoplasmic signal. Broken circle, NONO signal in nucleolus (Nuc) or nucleoplasm (NP). (*D*) NONO ChIP using site-specific primers. (*E*) Pull-down assay displaying 32P-γ-ATP end-labeled (32P*) gapmers by autoradiography after IP with immobilised HA-NONO variants FL and ΔRRM1 and PAGE separation. Silver stain and immunoblot, loading controls; dashed line, background: a.u. arbitrary units. *, p-value <0.05; **, p-value <0.001; two-tailed t-test. Error bar, mean ±SD. Representative images are shown. n=number of biological replicates.

### DNA damage reduces NONO occupancy at protein-coding gene promoters and attenuates pre-mRNA synthesis

DSB signaling inhibits RNAPII activity, in particular close to transcriptional start sites (TSSs) and NONO stimulates pre-mRNA synthesis in unperturbed cells (Iannelli et al. 2017; Wei et al. 2021). Thus, we speculated that the etoposide-induced nucleolar re-localisation of NONO coincides with reduced RNAPII activity on broken chromatin. To test this, we performed proximity ligation assays (PLAs) for NONO and RNAPII or the elongation factor SPT5 and observed prominent PLA signals for both costainings in unperturbed cells, which were sensitive to etoposide treatment (Fig. 3A). Thus, we investigated if DNA damage alters NONO chromatin occupancy at protein-coding genes and performed CUT&RUN-seq with the NONO antibody. We observed prominent binding of NONO in a region from the TSS up to 1 kb downstream of the TSS, but not the TES of highly expressed genes, which was markedly reduced upon etoposide treatment and sensitive to NONO depletion (Fig. 3B,C, Supplemental Fig. S5A,B). We validated CUT&RUN-seq data by NONO ChIP assays and detected NONO occupancy downstream of the TSSs of *ACTB* and *CCNB1*, which was sensitive to etoposide treatment (Supplemental Fig. S5C). Next, we applied 4sU-seq to measure nascent RNA synthesis. We found that etoposide treatment reduced the bulk of pre-mRNA synthesis by ∼25% within the gene body of highly expressed genes (Fig. 3D, Supplemental Fig. S5D). The depletion of NONO *per se* reduced 4sU-seq reads to similar extent, but not further reduced by combining NONO depletion with etoposide incubation. Thus, etoposide treatment depletes NONO from the promoter-proximal region of some highly expressed genes to attenuate RNAPII activity.

**Figure 3.**
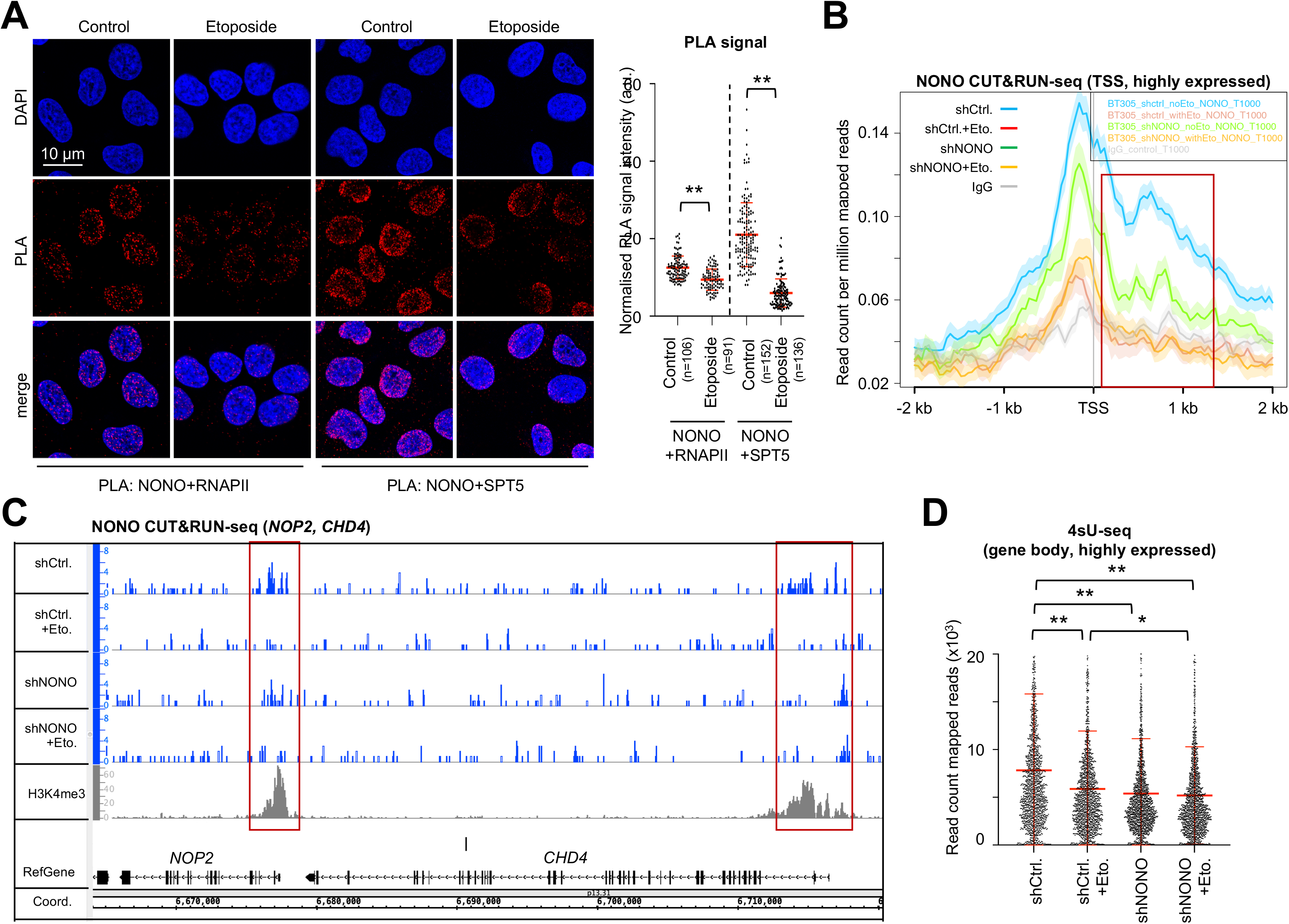
DNA damage reduces promoter-associated occupancy of NONO and RNAPII activity in U2OS cells. (*A*) Imaging (left) and quantitation (right) of proximity ligation assay (PLA) signals for NONO/RNAPII or NONO/SPT5. Each dot represents one acquisition. n, number of cells; a.u., arbitrary units. (*B*) NONO CUT&RUN-seq at the transcriptional start site (TSS) of top 1000 expressed genes. Red, promoter region. (*C*) Browser tracks of NONO and histone H3 lys-4 tri-methylation (H3K4me3) CUT&RUN-seq. Red, promoter region. (*D*) 4sU-seq read counts for the gene body of 863 highly expressed genes. *, p-value <0.05; **, p-value <0.001; two-tailed t-test. Error bar, mean ±SD. Representative images are shown. n=number of biological replicates.

### NONO mediates the accumulation of pre-mRNA transcripts in the nucleolus

NONO preferentially binds intron-containing transcripts in unperturbed cells (Xiao et al. 2019; Zhang et al. 2022). We reasoned that the DDR shifts NONO from protein-coding chromatin to the nucleolus to detain nascent transcripts from broken chromatin. To explore DNA damage-induced NONO-dependent changes in the nucleolar transcriptome, we created U2OS cells that stably express GFP-tagged ascorbate peroxidase 2 fused with three nucleolar targeting sequences from the NF-κB-inducing kinase (U2OS:GFP-APEX2-NIK3), which can be used to map nucleolar transcripts *in vivo* by proximity labeling and subsequent sequencing of immunoselected biotinylated RNA (APEX-seq) (Fazal et al. 2019) (Fig. 4A). We confirmed GFP-APEX2-NIK3-mediated biotinylation of nucleic acids by dot blotting (Supplemental Fig. S6A) and validated the selective biotinylation of RNA on agarose gels by immunoselection and RNaseA digestion (Supplemental Fig. S6B). We further confirmed nucleolar localisation and activity of the GFP-APEX2-NIK3 reporter by costaining with NCL and a fluorescently labeled neutravidin probe, irrespective of etoposide treatment (Supplemental Fig. S6C,D). Importantly, the expression of GFP-APEX2-NIK3 did not interfere with NONO nucleolar re-localisation, nor did the depletion of NONO interfere with GFP-APEX2-NIK3 localisation (Fig. 4B, Supplemental Fig. S6E). Reassuringly, we found that proximity-mediated biotinylation by nucleolar GFP-APEX2-NIK3 in the presence of etoposide increased the amount DBHS proteins, but not GFP-APEX2-NIK3 or fibrillarin, that coimmunoprecipitate with streptavidin beads 2-4-fold (Fig. 4C). This prompted us to perform APEX-seq in U2OS:GFP-APEX2-NIK3 cells. By assessing fold-changes for a total of 14463 intron-containing, biotin-labeled transcripts, we found 75 candidates with significantly higher biotinylation upon etoposide treatment (Fig. 4D). To exclude that the changes in the levels of biotinylated transcripts reflect lentiviral stress or perturbations upon biotin-phenol/H_2_O_2_ treatment, we compared the ratios of labeled transcripts from lentiviral transduced cells with rRNA-depleted transcripts immunoselected from unlabeled and unperturbed controls. We found no significant changes in the levels of biotinylated transcripts (Fig. 4E). To assess if the differential biotinylation of transcripts depends on NONO, we repeated APEX-seq upon NONO depletion. Strikingly, NONO depletion abolished the differential biotinylation, but not the synthesis of most candidates (Fig. 4F, Supplemental Fig. S6F). To assess if NONO binds candidates differentially upon DNA damage, we employed the NONO antibody for enhanced cross-linking immunoprecipitation and sequencing (eCLIP-seq). We found that etoposide increased the total number of NONO eCLIP-seq peaks from 2649 to 3991 and particularly enhanced NONO binding to intron-containing transcripts, including the previously identified transcript DAZAP1 (Zhang et al. 2022), and some of the identified APEX-seq candidates (CDKN1A, PURPL) (Fig. 4G,H, Supplemental Fig. S6G,H). As NONO binding to chromatin-associated transcripts is strongly correlated with the formation of R-loops (Wu et al. 2022), we tested if the defects in nucleolar detention observed in NONO-deficient cells may be linked to aberrant R-loop levels. We expressed the R-loop-stabilising V5-tagged RNaseH1 D210N mutant, or wild type control, and employed CUT&RUN-seq to assess R-loops globally. We found that the depletion of NONO prior to etoposide treatement increased the levels of R-loops within the gene body of APEX-seq candidates CDKN1A and BTG2 (Fig. 4I, Supplemental Fig. S6I). This suggests that NONO mediates nucleolar detention of pre-mRNA transcripts to mitigate R-loops level upon DNA damage.

**Figure 4.**
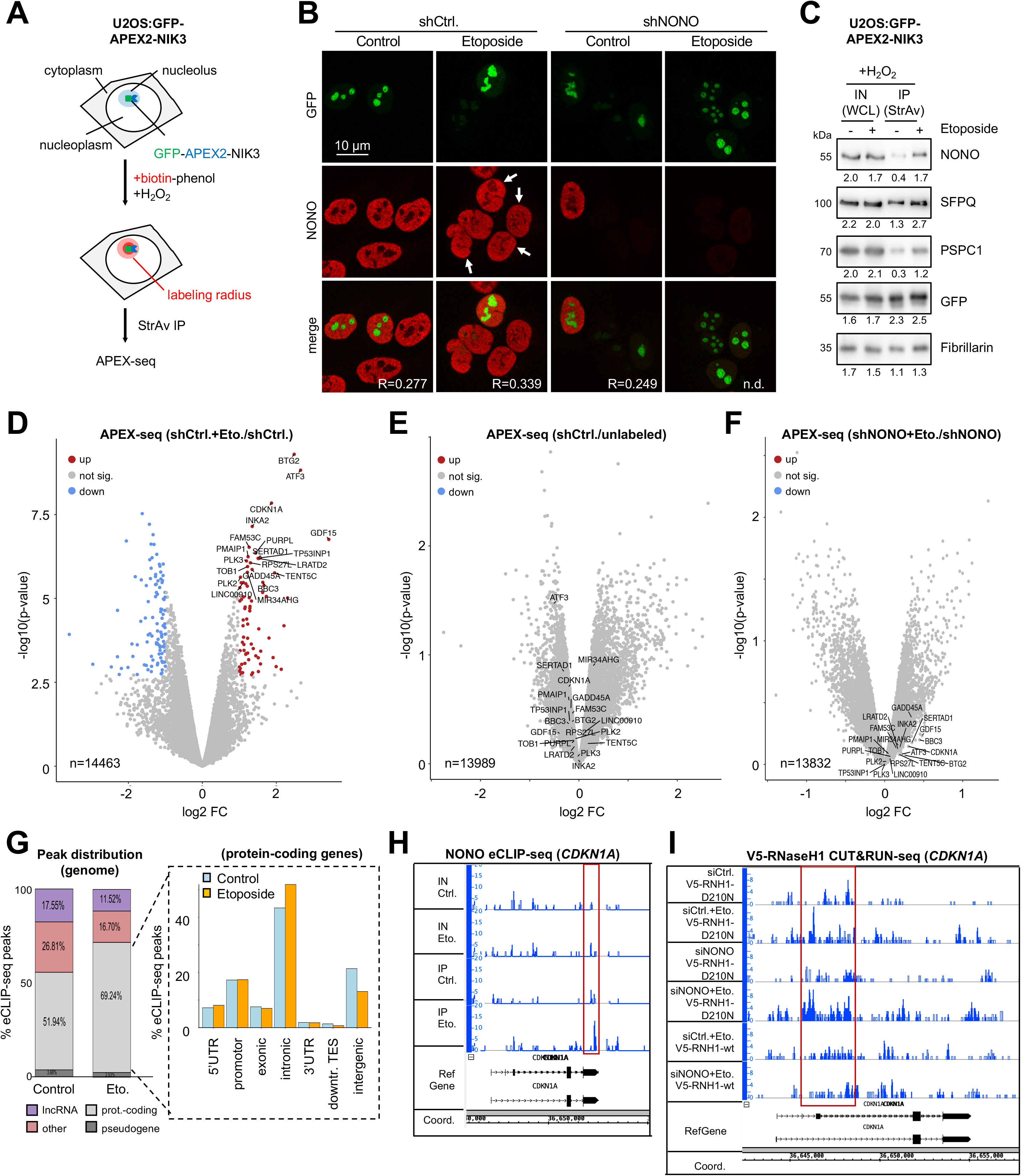
NONO mediates the nucleolar accumulation of transcripts in U2OS cells. (*A*) Schematic displaying APEX-seq in U2OS:GFP-APEX2-NIK3 cells. StrAv, streptavidin; H_2_O_2_, hydrogen peroxide. (*B*) Imaging of GFP and NONO in U2OS:GFP-APEX2-NIK3 cells. R=Pearson correlation; n.d., not detected; arrowhead, pan-nuclear NONO. Representative images are shown. (*C*) Immunoblots detecting NONO, SFPQ, PSPC1, GFP-APEX2-NIK3 and fibrillarin upon incubation with biotin-phenol, H_2_O_2_ from whole cell lysates (WCL) or upon immunoselection with streptavidin-coated beads. (*D-F*) Volcano plots displaying the relative abundance of transcripts as ratios of reads. Red, overrepresented; blue, underrepresented; n=number of transcripts. (*G*) NONO eCLIP-seq peak distribution genome-wide (left) and at the gene body (right). (*H*) Browser tracks for NONO eCLIP-seq reads. Red, increased binding. (*I*) Browser tracks depicting V5-RNaseH1 CUT&RUN-seq reads for *CDKN1A* ±NONO depletion/etoposide. Red box, region of increase.

### NONO inactivation impairs DSB signaling

NONO depletion elevates R-loop levels at telomeres and promotes genome instability (Petti et al. 2019). To test the impact of NONO depletion on DSB signaling, we performed etoposide incubation kinetics and detected defects in clearing ser-1981-phosphorylated (p)ATM and γH2A.X upon NONO depletion (Supplemental Fig. S7A). Complementation with mCherry-NONO rescued γH2A.X levels partially (Supplemental Fig. S7B). To investigate if the DDR function of NONO may be linked to its nucleolar re-localisation, we assessed the impact of RRM1 depletion on DSB signaling. We found that overexpression of ΔRRM1, but not FL, increased phosphorylation of DDR markers (Fig. 5A). Next, we asked if NONO depletion elevates the amount of DSBs and used breaks labeling *in situ* and sequencing (BLISS-seq) to quantify DSBs (Fig. 5B). Indeed, NONO depletion prior to etoposide treatment increased the amount of DSBs compared to non-depleted, etoposide-treated cells at TSSs of highly expressed genes. For validation, we performed γH2A.X ChIP at the AsiSI-site DS1 (Supplemental Fig. S7C). Again, NONO depletion prior to 4-OHT incubation increased γH2A.X levels about 2-fold. Interestingly, histone H2B acetylation at lys-120 residues (H2BK120ac) functions as chromatin switch during DSBR at AsiSI sites (Clouaire et al. 2018). Thus, we applied CUT&RUN-seq to quantify the levels of H2BK120ac at TSSs of highly expressed genes (Fig. 5C, Supplemental Fig. S7D). The depletion of NONO or etoposide treatment alone modestly altered H2BK120ac levels at TSSs. Combining NONO depletion with etoposide treatment, however, strongly increased the H2BK120ac mark. Finally, we rescued elevated H2BK120ac levels by reexpression of mCherry-NONO (Supplemental Fig. S7E). We conclude that NONO inactivation impairs DSB signaling.

**Figure 5.**
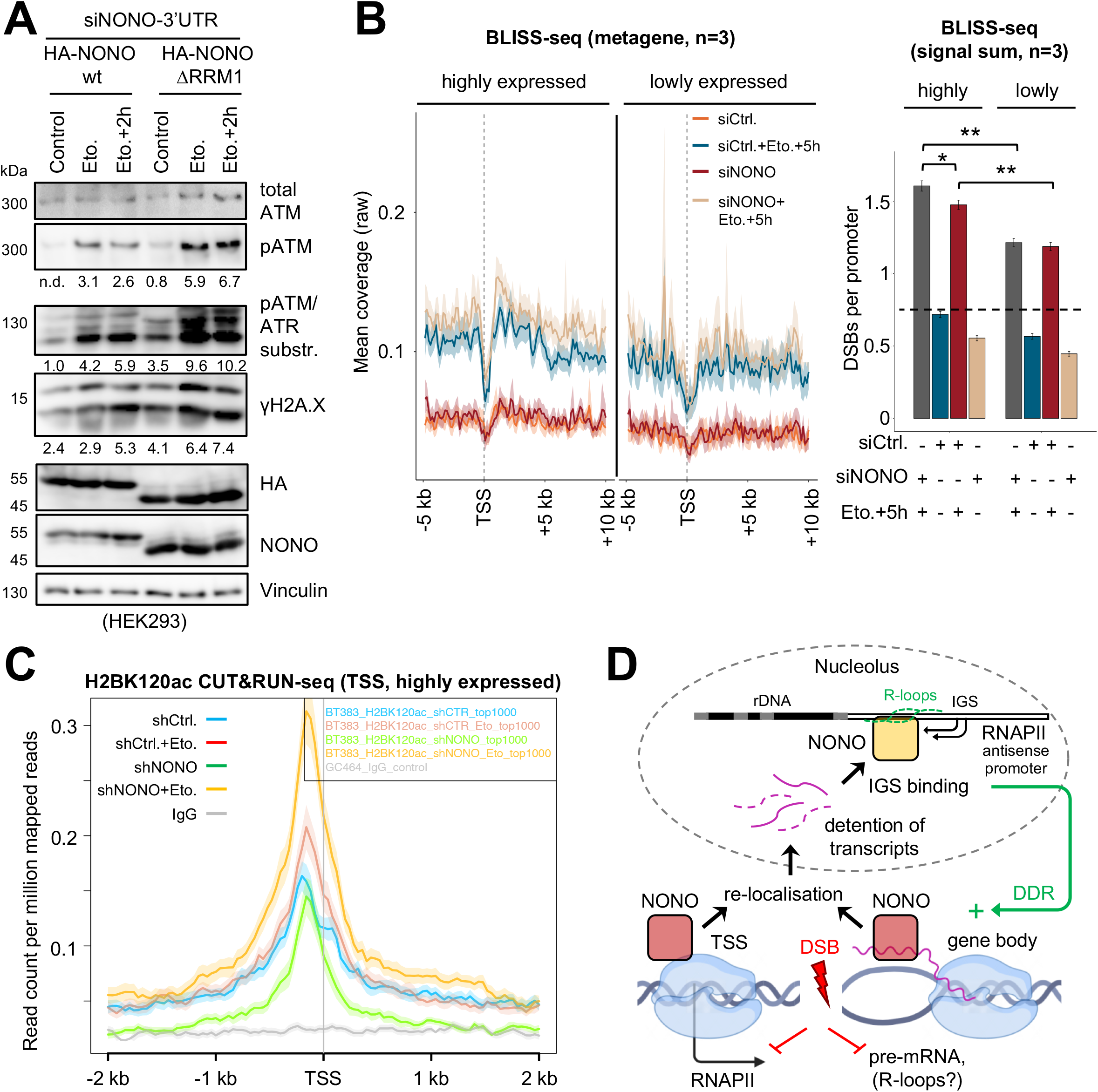
Impairment of NONO interferes with DSB signaling in U2OS cells. (*A*) Immunoblots detecting total ATM, pATM, pATM/ATR substrates, γH2A.X, HA-NONO and endogenous NONO. (*B*) BLISS-seq metagene profiles (left) and signal sum (right) detecting DSBs at the TSS of highly and lowly expressed genes. Dashed line, background. (*C*) H2BK120ac CUT&RUN-seq at TSSs of top 1000 expressed genes. *, p-value <0.05; **, p-value <0.001; two-tailed t-test. Error bar, mean ±SD. Representative images are shown. n=number of biological replicates. (*D*) Model illustrating our findings.

## Discussion

We describe NONO as attenuator of pre-mRNA synthesis and nucleolar detainer of nascent transcripts to promote DSBR (Fig. 5D). Many RBPs display stress-induced nucleolar re-localisation (Mamontova et al. 2021; Feng and Manley 2022). We provide evidence for diNAR-induced nucleolar R-loops as anchor for nucleolar NONO. How is diNAR synthesis regulated? IGS loci may become accessible for RNAPII upon DNA damage-induced looping of nucleolar DNA to the nucleoplasm. Alternatively, RNAPII elongation factors may enrich in the nucleolus, as shown for Spt4 in yeast (Yokoyama et al. 2023). Inhibition of RNAPI may also enhance diNAR synthesis. DSB signaling indeed attenuates RNAPI transcription via ATM when DSBs occur both in the nucleolus and the nucleoplasm (Korsholm et al. 2020; Li and Yan 2023). However, we did not observe NONO nucleolar re-localisation upon induction of nucleolar DSBs with I-PpoI. This suggests that the initial DNA-damaging events that trigger NONO re-localisation occur in the nucleoplasm, but may also involve attenuation of RNAPI activity.

We observed a rapid decrease in NONO chromatin occupancy downstream of some protein-coding gene promoters within 2 hours of etoposide treatment, whilst NONO accumulation at nucleolar IGS loci was prominently detected upon chase. Thus, the reduction of NONO at promoter regions likely precedes its re-localisation to nucleoli and impairs pre-mRNA synthesis as a consequence thereof. Early studies identified NONO as transducer of cAMP signaling that interacts with the CBP/p300 coactivator complex, copurifies with the mediator complex and associates with the RNAPII CTD (Yang et al. 1997; Emili et al. 2002; Amelio et al. 2007). NONO enriches in condensates to enhance the expression of pre-mRNA transcripts in neuroblastoma (Zhang et al. 2022). Other DBHS proteins also stabilise nascent pre-mRNA and favour the placement of RNAPII-activating CTD phospho-marks at promoters (Shao et al. 2022). This suggests that DBHS proteins foster RNAPII activity and that NONO nucleolar re-localisation diminishes RNAPII-stimulating conditions.

NONO nucleolar re-localisation attenuates pre-mRNA synthesis to mitigate aberrant transcripts via nucleolar shielding. This could promote R-loop-dependent DSBR pathway choice. R-loops accumulate at actively transcribed DSBs (Bader and Bushell 2020). R-loops foster the recruitment of critical homologous recombination factors to DSBs (Hatchi et al. 2015; D’Alessandro et al. 2018). R-loops also promote DNA end resection via DNA endonuclease CtIP, a critical step in DSBR pathway choice (Gómez-Cabello et al. 2022). Thus, the NONO-mediated nucleolar detention of transcripts may suppress R-loops and, at least in part, explain promotion of NHEJ by NONO (Krietsch et al. 2012; Jaafar et al. 2017). Overall, we provide evidence for a nucleolar DDR that engages NONO to shield aberrant transcripts from DSBs.

## Materials and methods

### Tissue culture

Human U2OS, AsiSI-ER expressing U2OS (gift from Gaelle Legube), GFP-APEX2-NIK3 expressing U2OS (U2OS:GFP-APEX2-NIK3) and HEK293 cells were cultured in Dulbecco’s modified eagle’s medium (DMEM, Gibco) with 10% fetal bovine serum (FBS, Capricorn), 100 U/mL penicillin-streptomycin (Gibco), 2 mM L-glutamine (Gibco) at 37°C and 5% CO_2_. Cells were incubated with etoposide (Sigma, 20 µM), THZ1 (Biozol, 1 µM), CX-5461 (Selleckchem, 1 µM), for 2 h, 4-OHT (Sigma, 10 µM) for 4 h, unless stated differently.

### Transfection and viral work

Transfection of expression plasmids (Supplemental Table S1) was performed with Lipofectamine 2000 (Invitrogen) and Opti-MEM (Gibco) using the manufacturer’s protocol. HA-NONO mutants were PCR-cloned with primers (Supplemental Table S2) and a Q5 site-directed mutagenesis kit (NEB) using the manufacturer’s protocol and verified by sequencing. siRNA (100 nM) was transfected (6 h) on two consecutive days. Short-hairpin (sh)RNA were transduced by lentiviral infection (Supplemental Table S2). To generate U2OS:GFP-APEX2-NIK3 cells, 10 µg pLX304-GFP-APEX2-NIK3 plasmid (gift from Alice Ting) was pooled with psPAX2 and pMD2.G (gift from Elmar Wolf), mixed with 30 µL polyethylenimine (Calbiochem), mixed in 500 µL OptiMEM, incubated (25 min, RT), added to HEK293 cells preincubated in 5 mL DMEM/2% FBS, and transfected (8 h). Virus was harvested, sterile filtered and frozen. U2OS cells were infected (24 h) in viral mixture (1.5 mL DMEM, 1.5 mL viral harvest, 6 µL polybrene, Invitrogen) and DMEM with 7.5 µg/mL blasticidin (Sigma) for polyclonal selection (10 days).

### Protein analytics

Proteins were assessed as whole cell extracts, directly lysed, boiled and sonicated in 4x sample buffer (250 mM tris-HCl pH6.8, 8% SDS, 40% glycerol, 8% β-mercaptoethanol, 0.02% bromophenol blue). Samples were separated by SDS-PAGE and stained with a SilverQuest kit (Invitrogen) or transferred to nitrocellulose membranes (Cytivia), blocked and washed in PBS/0.1% triton x-100/5% milk (PBST), probed with selective antibodies (Supplemental Table S3) and visualised with an ECL kit (Cytivia) on an imaging station (LAS-4000, Fuji). Signals were quantified by ImageJ (NIH). For immunoprecipitation (IP), cells were trypsinised, washed in PBS and centrifuged (1200 rpm, 5 min). Pellets were lysed (10 min on ice) in 5 volumes IP buffer (200 mM NaCl, 0.5 mM EDTA, 20 mM HEPES, 0.2% NP-40, 10% glycerol, 400U Ribolock inhibitor, 1x protease/phosphatase inhibitor, Roche). Lysates were centrifuged (12000 rpm, 12 min) and supernatants were incubated (2 h, 4°C) with 5 µg antibodies conjugated to 25 µL protein G dynabeads (Invitrogen). Samples were immobilised, washed in IP buffer (10 min, 4°C) and eluted with sample buffer (5 min, 95°C).

### Pull-down assays

100 pmol gapmers (Supplemental Table S4) were labeled with biotin-16-ddUTP (Jena) and a 2^nd^ generation DIG-oligonucleotide 3’end-labeling kit (Roche) using the manufacturer’s protocol or with radioactive labeling mix (1 µL 10x PNK buffer, NEB, 1 µL of 100 µM gapmer, 1 µL T4 PNK, NEB, 1 µL γ-^32^P-ATP, Hartmann, 6 µL ddH_2_O) for 40 min at 37°C. End-labeled gapmers were centrifuged (3200 rpm, 5 min) with G-25 columns (Cytivia), diluted in 800 µL IP buffer and incubated (2 h, RT with rotation) with either 0.4 µg recombinant NONO (rec-NONO) (ActiveMotif) or HA-NONO variants that were immobilised on HA-conjugated protein G dynabeads upon expression in HEK293 cells and IP. Rec-NONO complexes were captured on 25 µL streptavidin C1 dynabeads (Invitrogen), washed in IP buffer, eluted by boiling (95°C, 5 min) in sample buffer and analysed by immunoblotting. HA-NONO complexes were washed in IP buffer, split and either eluted as above or by heating (65°C, 5 min) in 2x loading dye (7 M urea, 0.05% xylene cyanol, 0.05% bromophenol blue) for separation by UREA-PAGE (30 min, 350 V) in 1x TBE buffer (90 mM tris, 90 mM boric acid, 2 mM EDTA), transfer on whatman paper with a gel-dryer (BioRad) and detection by autoradiography and films (Cytivia).

### BLISS-seq

Cells were washed in PBS, fixed (10 min, RT) with 5% paraformaldehyde, washed with PBS, lysed (1 h, 4°C) in lysis buffer 1 (10 mM tris-HCl pH8.0, 10 mM NaCl, 1 mM EDTA, 0.2% triton x-100), washed in PBS, lysed again (1 h, 37°C) in lysis buffer 2 (10 mM tris-HCl pH8.0, 150 mM NaCl, 1 mM EDTA, 0.3% SDS) and washed in PBS. For AsiSI digestion, samples were equilibrated (2 min, RT) in 150 µL CutSmart buffer (NEB). 3µl recombinant AsiSI endonuclease (10U/µL, NEB) was added to three biological replicates and incubated (2 h, 37°C). Controls were incubated in buffer only. For DSB blunting, samples were washed in CutSmart buffer, and incubated (1h, RT) in 150 µL blunting mix (112.5 µL ddH_2_O, 15 µL 10x blunting buffer, NEB, 15 µL 100 µM dNTPs, 0.3 µL 50 mg/mL BSA, 6 µL blunting enzyme mix from quick blunting kit, NEB). Prior to ligation, 10 µM of corresponding BLISS adapters (Supplemental Table S5) were mixed equimolar and annealed (5 min, 95°C with gradient cooling to 25°C). For ligation, samples were washed in CutSmart buffer, preincubated (5 min, RT) in 1x T4 ligase buffer (NEB), and incubated (18 h, 16°C with gentle shaking) in 150 µL ligation buffer (124.5 µL ddH_2_O, 15 µL 10x T4 ligase buffer, 3 µL 50 mg/mL BSA, 1.5 µL 2000U/µL T4 ligase, NEB, 6 µL BLISS adapter pairs). For removal of excess adapters, samples were incubated (1 h, 37°C, with gentle shaking) in high salt wash buffer (10 mM tris-HCl pH8.0, 2 M NaCl, 2 mM EDTA, 0.5% triton x-100) and washed in PBS. For extraction of genomic DNA, samples were incubated (5 min, RT) in 100 µL extraction buffer (10 mM tris-HCl pH8.0, 100 mM NaCl, 50 mM EDTA, 1% SDS, 10% 10 mg/mL proteinase K, Sigma), harvested by scaping, pooled (merged conditions for each replicate), and incubated (18 h, 55°C). DNA was purified by phenol/chloroform extraction, recovered in 50 µL ddH_2_O, and sonicated (Covaris). Fragmented DNA was concentrated with SPRI select beads (Beckman) and a magnet (Alpaqua), washed with 80% ethanol, air-dried and eluted in 8 µL ddH_2_O. For in vitro transcription (IVT), 7.5 µL DNA was incubated (14 h, 37°C) with IVT mix (0.5 µL Ribolock inhibitor, Invitrogen, 2 µL T7 polymerase buffer, NEB, 8 µL rNTP mix, 2 µL T7 polymerase, Invitrogen). DNA was removed by addition of 1 µL turbo DNase (Invitrogen) for 15 min. RNA size selection and clean-up was performed with RNAClean XP beads (Beckman), size selected RNA was washed in 80% ethanol, air-dried and eluted in 6 µL ddH_2_O. For library preparation, 1 µL of 5 µM RA3 adapter (NEB) was added to 5 µL RNA sample, incubated (2 min, 70°C) and placed on ice. 4 µL ligation mix (2 µL 10x T4 ligase buffer, NEB, 1 µL T4 RNA ligase 2, truncated, NEB, 1 µL Ribolock inhibitor) was added and incubated (1 h, 28°C). For reverse transcription (RT), 3.5 µL ddH_2_O and 1 µL 10 µM RTP primer (NEB) was added, incubated (2 min, 70°C) and placed on ice. 5.5 µL RT mix (2 µL 5xGC buffer, Invitrogen, 0.5 µL 12.5 mM dNTP mix, 1 µL 100 mM DTT, 1 µL SuperScriptIII reverse transcriptase, Invitrogen, 1 µL Ribolock inhibitor) was added and incubated (1 h, 50°C) and heat inactivated (15 min, 70°C). For indexing and amplification, 10 µL of RT reaction was mixed with 25 µL NEBNext 2x PCR mix, 2 µL 10 µM RPI primer (NEB), 2 µL 10 µM RP1 primer (NEB), 1 µL ddH_2_O and PCR amplified for 16-18 cycles. Library clean-up, was performed with AMPure XP beads (Beckman). The library was captured, washed with 80% ethanol, air-dried, eluted in 20 µL ddH_2_O prior to sequencing.

### ChIP and CUT&RUN-seq

For ChIP, cells were fixed with 1% formaldehyde (10 min, 37°C), quenched in 0.125 M glycine (10 min, 37°C), washed in PBS and centrifuged (2000 rpm, 5 min). Pellets were resuspended in 500 µL cold cell lysis buffer (5 mM PIPES pH8.0, 85 mM KCl, 0.5% NP-40, 1x protease/phosphatase inhibitor) and lysed (10 min on ice). Nuclei were centrifuged (3000 rpm, 5 min) and resuspended in 400 µL cold nuclear lysis buffer (1% SDS, 10 mM EDTA, 50 mM tris-HCl pH8.0, 1x protease/phosphatase inhibitor) and lysed (10 min on ice). Lysates were sonicated (5x 5 min, 30 sec on/off) with a Bioruptor (Diagenode) and pelleted (13000 rpm, 10 min). The supernatant was mixed with 2 mL dilution buffer (0.01% SDS, 1.1% triton x-100, 1.2 mM EDTA, 16.7 mM tris-HCl pH8.0, 167 mM NaCl, 1x protease/phosphatase inhibitor). Diluted samples were aliquoted, 5 µg antibodies were added (IP sample) or not (input) and incubated overnight (4°C with rotation). For pull-down, 20 µL of protein G dynabeads were added to IP samples, incubated (1.5 h with rotation), immobilised and washed in wash buffer A (0.1% SDS, 1% triton x-100, 2 mM EDTA, 20 mM tris-HCl pH8.0, 150 mM NaCl), B (0.1% SDS, 1% triton x-100, 2 mM EDTA, 20 mM tris-HCl pH8.0, 500 mM NaCl), C (0.25 M LiCl, 1% NP-40, 1% sodium deoxycholate, 1 mM EDTA and 10 mM tris-HCl pH8.0), and twice with D (10 mM Tris-HCl pH8.0, 1 mM EDTA). For elution, samples were incubated with 500 µL elution buffer (1% SDS, 0.1 M NaHCO_3_) for 30 min with rotation. Reversal of cross-links was performed at 65°C overnight after adding 30 µL 5 M NaCl, 1 µL 10 µg/mL RNaseA (Sigma), 10 µL 0.5 M EDTA, 20 µL 1 M tris-HCl pH6.5, 2 µL 10 mg/mL proteinase K (Sigma) to input and IP samples. DNA was purified by phenol/chloroform extraction, recovered in ddH_2_O, assessed by qPCR with selective primers (Supplemental Table S6).

For DNA-RNA hybrid IP (DRIP) non-crosslinked lysates were incubated (1 h, 37°C) with 10U RNaseH (NEB) prior to immunoselection. For CUT&RUN-seq, cells were harvested with accutase (Sigma), centrifuged (600 rpm, 3 min) and washed in wash buffer (20 mM HEPES pH7.5, 150 mM NaCl, 0.5 mM spermidine). Cells were incubated (10 min, RT) with 10 µL concanavalinA-coated magnetic beads (BioMag) resuspended in an equal volume of binding buffer (20 mM HEPES pH7.5, 10 mM KCl, 1 mM CaCl_2_, 1 mM MnCl_2_), immobilised, permeabilised with 150 µL antibody buffer (20 mM HEPES pH7.5, 150 mM NaCl, 0.5 mM spermidine, 0.05% digitonin, 2 mM EDTA) and incubated with 1 µg primary antibody (800 rpm, 4°C, overnight with rotation). Samples were immobilised, washed in dig-wash buffer (20 mM HEPES pH7.5, 150 mM NaCl, 0.5 mM spermidine, 0.05% digitonin) and incubated (1 h, 800 rpm, 4°C with rotation) with 150 µL protein A/G-MNase fusion protein (1 µg/mL, CST). Samples were immobilised, washed in dig-wash buffer and once with 1 mL rinse buffer (20 mM HEPES pH7.5, 0.05% digitonin, 0.5 mM spermidine). For chromatin digestion and release, samples were incubated (30 min, on ice) in cold digestion buffer (3.5 mM HEPES pH7.5, 10 mM CaCl_2_, 0.05% digitonin). The reaction was stopped by addition of 200 µL stop buffer (170 mM NaCl, 20 mM EGTA, 0.05% digitonin, 50 µg/mL RNaseA, 25 µg/mL glycogen) and fragments were released by incubation (30 min, 37°C). The supernatant was incubated (1 h, 50°C) with 2 µL 10% SDS and 5 µL proteinase K (10 mg/mL, Sigma). Chromatin was recovered by phenol/chloroform extraction and resuspended in 30 µL TE (1 mM tris-HCl pH8.0, 0.1 mM EDTA). For sequencing, three biological replicates were quantified with a fragment analyser (Advanced Analytical), pooled and subjected to library preparation. Libraries for small DNA fragments (25-75 bp) were prepared with NEBNext Ultra II DNA library prep Kit (NEB#E7645).

### RNA analytics

Total RNA was isolated by TRIzol (Invitrogen) using the manufacture’s protocol. cDNA was synthesised with SuperScriptIII enzyme (Invitrogen) and gene-specific primers (Supplemental Table S6) and quantified upon reverse transcription quantitative PCR (RT-qPCR) with PowerUp SYBR green master mix (Applied) using the manufacturer’s protocol. For dot blots, total RNA was extracted by TRIzol, resuspended in ddH_2_O with 0.02% methylene blue, heated (5 min, 72°C), spotted on a nylon membrane (Cytivia), crosslinked (120 mJ/cm^2^) in a crosslinker (UVP), blocked in PBS/0.1% triton x-100/0.5% SDS (20 min), washed in PBS/0.1% triton x-100 (20 min), incubated (4°C, overnight) with a streptavidin-HRP probe (Invitrogen), washed in PBS/0.1% triton x-100 (20 min), and visualised with an ECL kit (Cytivia). For SYBR gold (Invitrogen) staining, immunoselected transcripts were on-bead digested (10 min, RT) with 2 µL 10 µg/mL RNaseA (Sigma), separated by UREA-PAGE, stained with 1x SYBR gold diluted in 1x TBE (10 min in the dark) and visualised on a transilluminator (Thermo).

### mNET-seq

For mNET-IP, 5 µg antibodies were coupled to protein G dynabeads, washed and resuspended in 100 µL NET-2 buffer (50 mM tris-HCl pH7.4, 150 mM NaCl, 0.05% NP-40). Cells were harvested, washed in PBS and lysed in hypotonic buffer (10 mM HEPES pH7.9, 60 mM KCl, 1.5 mM MgCl_2_, 1 mM EDTA, 1 mM DTT, 0.075% NP-40, 400U Ribolock inhibitor, 1x protease/phosphatase inhibitor) (10 min, 4°C with rotation). Nuclei were centrifuged (2 min, 1000 rpm), washed in hypotonic buffer without NP-40 and resuspended in 125 µL cold NUN1 buffer (20 mM tris-HCl pH7.9, 75 mM NaCl, 0.5 mM EDTA, 50% glycerol, 400U Ribolock inhibitor, 1x protease/phosphatase inhibitor). 1.2 mL NUN2 buffer (20 mM HEPES-KOH pH7.6, 300 mM NaCl, 0.2 mM EDTA, 7.5 mM MgCl_2_, 1% NP-40, 1 M urea, 400U Ribolock inhibitor, 1x protease/phosphatase inhibitor) was added and nuclei were incubated (on ice, 15 min) and centrifuged (10 min, 16000 rpm). Non-soluble chromatin pellet was washed in 100 µL 1x MNase buffer (NEB), centrifuged and digested (2 min, 37°C with rotation) in 100 µL MNase reaction mix (87 µL ddH_2_O, 10 µL 10x MNase buffer, NEB, 1 µL 100x BSA, 2 µL 2000 U/µL MNase, NEB). Digests were centrifuged (5 min, 16000 rpm) and the supernatant was diluted with 10 volumes NET-2 buffer. Conjugated antibodies were added and incubated (2 h, 4°C with rotation). Samples were immobilised and washed in NET-2 buffer. For analysis of proteins, input and mNET-IP samples were analysed by immunoblotting as above. For analysis of transcripts, 10% of mNET-IP sample was subjected to TRIzol extraction and RT-qPCR or end-labeled on beads with radioactive PNK labeling mix and analysed by autoradiography or monitored for enrichment by immunoblotting. 90% of mNET-IP sample was end-labeled on beads with non-radioactive PNK labeling mix, eluted and separated by UREA-PAGE along with inputs. A small RNA (<100 nts) fraction was size-selected according to methylene blue migration. Slices were incubated (2 h, RT with rotation) in 400 µL elution buffer (1 M NaOAc, 1 mM EDTA), centrifuged (2 min, 13000 rpm). Supernatants containing eluted RNA were loaded on spin-x-columns (Coster) and centrifuged (1 min, 13000 rpm). Flow-through was precipitated with 1 mL 100% ethanol and 1µL glycogen (Invitrogen), incubated (20 min, RT) and centrifuged (20 min, 13000 rpm). Pellets were washed in 70% ethanol, air-dried and recovered in 6 µL ddH_2_O. Three biological replicates were pooled and subjected to library preparation. Libraries were prepared with NEBNext Multiplex small RNA library prep Kit (NEB#7300) using the manufacturer’s protocol.

### 4sU-seq and APEX-seq

For 4sU-tagging, cells were incubated with 4sU (Sigma, 2 mM) for 15 min, directly lysed in 2.1 mL QIAzol (Qiagen), spiked with 4sU-labeled mouse cell lysates, and total RNA was extracted with miRNeasy kit (Qiagen) using the manufacturer’s protocol. 50 µg total RNA were diluted in 100 µL ddH_2_O, denaturated (5 min, 65°C), put on ice (10 min) and incubated (2 h, RT) with 50 µL biotin-HPDP (Thermo, 1.85 mM) diluted in 100 µL 2.5x biotin labeling buffer (25 mM tris-HCl pH7.4, 2.5 mM EDTA). The reaction was mixed with an equal volume of chloroform/isoamyl alcohol (24:1) and separated with a phase-lock tube (Qiagen) by centrifugation (14000 rpm, 5 min). RNA was precipitated (5 min, RT) with 1 µL glycogen (Invitrogen), 20 µL 5 M NaCl and an equal volume of isopropanol and centrifuged (14000 rpm, 20 min). The pellet was washed in an equal volume of 75% ethanol, centrifuged (14000 rpm, 10 min) and resuspended in 100 µL ddH_2_O. For APEX2-mediated proximity labeling, cells were incubated (30 min, 37°C) with 0.5 mM biotin-phenol (Iris), pulsed (1 min) with 1 mM H_2_O_2_ (Sigma) and quenched by 10 mM sodium ascorbate (Sigma), 5 mM trolox (Sigma) and 10 mM sodium azide (Sigma). Cells were directly lysed in 2.1 mL QIAzol (Qiagen) and total RNA was extracted with miRNeasy kit (Qiagen). For selection of biotinylated transcripts, samples were incubated (15 min, RT) with 50 µL streptavidin T1 dynabeads (Thermo), resuspended in an equal volume of 2x washing buffer (2 M NaCl, 10 mM tris-HCl pH7.5, 1 mM EDTA, 0.1% tween-20). Samples were immobilised and washed in washing buffer (1 M NaCl, 5 mM tris-HCl pH7.5, 0.5 mM EDTA, 0.05% tween-20). For elution, samples were incubated with 100 µL DTT (100 mM) at RT for 5 min and recovered by the RNeasy clean up kit (Qiagen) using the manufacturer’s protocol. For APEX-seq, biotinylated RNA was enriched by incubation with 20 µL streptavidin C1 dynabeads (2h, 4°C) washed in washing buffer (5 mM tris-HCl, pH 7.5, 0.5 mM EDTA, 1 M NaCl; 0.1% TWEEN 20) and solution A (100 mM NaOH, 50 mM NaCl) and resuspended in solution B (100 mM NaCl). Samples were immobilised and washed in washing buffer and resuspended in 54 µL ddH_2_O. For elution, samples were incubated (1 h, 42°C followed by 1 h, 55°C) with 54 µL 3x proteinase digestion buffer (330 µL 10x PBS, 330 µL 20% N-laurylsarcosine sodium solution, 66 µL of 0.5 M EDTA, 16.5 µL of 1 M DTT, 357.5 µL ddH_2_O, 10 µL proteinase K, 2 µL Ribolock) and recovered by RNA clean and concentrator kit (Zymo) using the manufactureŕs protocol. For 4sU-sequencing and APEX-seq, samples were quantified by RiboGreen assay (Thermo) using the manufacturer’s protocol, subjected to library preparation as individual replicates (4sU-seq) or pooled replicates (APEX-seq). Libraries were prepared with NEBNext Ultra II Directional RNA library prep Kit (NEB#E7760) and NEBNext rRNA Depletion Kit (NEB#E6310) using the manufacturer’s protocol.

### eCLIP-seq

Cells cultured in the absence or presence of etoposide were washed in PBS, subjected to UV irradiation (200 mJ/cm^2^), scraped, resuspended in cold PBS, pelleted (1200 rpm, 5 min) and stored at −80°C. The pellets were lysed (20 min, 4°C) in eCLIP lysis buffer (50 mM tris-HCl pH7.4, 150 mM NaCl, 1 mM EDTA, 1% NP-40, 0.5% sodium deoxycholate, 0.25 mM TCEP). After limited RNaseI (Invitrogen) and TURBO DNase (Invitrogen) digestion (20 min, 37°C), 5 µg NONO antibody was coupled to 30 µL protein G dynabeads and incubated with lysates (4°C, overnight). The samples were washed in eCLIP lysis buffer, in wash buffer (50 mM tris-HCl pH7.4, 300 mM NaCl, 1 mM EDTA, 1% NP-40, 0.5% sodium deoxycholate, 0.25 mM TCEP), followed by two washes in low-salt wash buffer (50 mM tris-HCl pH7.4, 1 mM EDTA, 0.5% NP-40). Subsequent library preparation was performed as described (Van Nostrand 2016).

### Imaging

Cells grown on cover slips (Roth) were washed in PBS, fixed (10 min) in 3% paraformaldehyde (Sigma), washed in PBS, permeabilised with PBS/0.1% triton x-100 (10 min) and blocked with PBS/10% FBS (2 h, 4°C). Primary and secondary antibodies (Supplemental Table S3) were diluted in PBS/0.15% FBS and incubated in a humidified chamber (overnight, 4°C or 2 h, RT), respectively. Cells were washed between incubations with PBS/0.1% triton x-100, sealed in 6-diamidino-2-phenylindole (DAPI)-containing mounting medium (Vectashield), and imaged by confocal microscopy (Leica-SP2, 63x, airy=1, sequential acquisition between frames, equal exposure times). Pan-nuclear localisation was scored in cells that display homogenous nuclear staining and colocalisation with nucleolar markers based on RGB profiler and Pearson’s correlation coefficient (ImageJ). PLAs were performed with a Duolink in-situ PLA kit (Sigma) using the manufacturer’s protocol. RNA-FISH experiments employ 30 non-overlapping, Quasar570-labeled sense DNA probes reverse complementary to 0.6 kb of the mapped region of antisense transcription at IGS-22 (Stellaris probe designer, masking level ≥2, Biosearch, Supplemental Table S7) using the manufacturer’s protocol.

### Statistics and bioinformatics

For APEX-seq, CUT&RUN-seq and 4sU-seq, base calling was performed with Illumina’s FASTQ Generation software v1.0.0 and quality was tested by FastQC. Reads were mapped with STAR (4sU-seq) (Dobin et al. 2013) or Bowtie2 (Langmead and Salzberg 2012) (other) to human hg19, human T2T, mouse mm10 or *E.coli* genome. Mouse reads for spike-normalisation were used as described (Orlando et al. 2014). CUT&RUN-seq read normalisation was performed by the sample-wise division of hg19-mapped reads by *E.coli*-mapped reads or read depth. The ratio was multiplied with the smallest number of *E.coli*-mapped reads. For 4sU-seq, reads falling in introns were considered, spike-normalised, sorted and indexed with SAMtools. Bedgraph files were generated with the genomecov function from BEDTools (Quinlan and Hall 2010). Density files were visualised by Integrated Genome Browser (IGB).

CUT&RUN-seq density plots were generated with ngs.plot using normalised bam files, testing top 1000 expressed genes in U2OS (Lorenzin et al. 2016). 1% extreme values were trimmed (option “–RB 0.01”). For APEX-seq, gene expression was assessed with featureCounts (Liao et al. 2014) on bam files with intron-containing, non-spliced reads. Differential gene expression was assessed with edgeR (Galaxy) using Benjamini Hochberg p-value <0.05, rejecting genes with <100 counts, and excluding non/weakly expressed genes. For 4sU-seq, counts that overlap between the spike normalised bam files and Human Genes (GRCh37.p13) were assessed with bedtool Intersect intervals (Galaxy), testing top 1000 expressed genes in U2OS (Lorenzin et al. 2016). The read count mapped reads for each condition were used for scatter plots in GraphPad.

For mapping of mNET-seq data to rDNA loci, FASTQ files were aligned to a custom reference genome based on U13369.1. Alignment was performed with bowtie2 allowing 1 mismatch and aligned reads were normalised to the sample with minimum aligned reads. Aligned reads were converted into bedGraphs carrying equal number of reads in the rDNA region and visualised by IGB. For analysis of IGS loci, paired-end samples were mapped to human genome CHM13 (version 1.1) with bowtie2 an preset parameter “very-sensitive-local”, and normalised to spiked-in reads mapping to mm10. A bed file with the coordinates of 214 rDNA-IGS (arranged in 5 clusters) was extracted from the gff3-file for CHM13 draft annotation v1.1, and used to generate a multifasta file with 6.8 million nucleotides. 470 *Alu* repeat sequences (representing 50 sub-families) were derived from hg38, with coordinates from the rmsk-table at UCSC (total length 112.000bp). IGS sequences not matching *Alu* elements were identified with blastn, resulting in 6.3 million nucleotides of “non-*Alu*” IGS, which were divided into bins of 100 nt. The number of spike-normalised ChIPseq reads mapping to each bin was determined with bedtools intersect. Numbers for bins overlapping individual IGSs were pooled.

BLISS-seq samples were demultiplexed based on their condition-specific barcodes with UMI-tools, allowing 1 mismatch, separately mapped to hg19 using Bowtie2 (default parameters) and filtered against an ENCODE Blacklist file to remove regions of high variance with bedtools intersect. For quantification of DSBs duplicated reads were identified by UMI, grouped and deduplicated with UMI-tools (default parameters). Density profiles were generated by R (package metagene2, assay parameter ‘ChIPseq’, 200 bp read extension). Bar graph was generated with by R (package exomeCopy) in the respective regions up- and downstream of the annotated TSS and divided by the number of genes in the corresponding gene set. Publicly available RNA-seq data (ENCODE: ENCFF182XEY) were filtered by gene length (≥1500 bp) to stratify genes by expression into highly (FPKM >10) and lowly (FPKM≤1) expressed genes. Promoters with proximal downstream TSSs were removed.

Paired-end sequencing reads from eCLIP experiments were trimmed with a custom Python script to identify the UMI and aligned to hg38 with the Burrows–Wheeler Aligner (BWA). PCR duplicates were removed by Picard’s MarkDuplicates with UMI-aware deduplication. Enriched protein-binding regions were identified by MACS2 callpeak (parameters ‘-g hs -s 58 -B --keep-dup all --nomodel --extsize 50 --d-min 5 --scale-to small –B’, comparing IP and size-matched IN samples). Visualisations of regions were rendered from the PCR-deduplicated.bam files by IGB. Distribution analysis employed ChIPpeakAnno package and TxDb.Hsapiens.UCSC.hg38.knownGene dataset, with plot generated using ggplot2.

## Data availability

NGS data are available at the gene expression omnibus (accession number GSE233594).

## Acknowledgments

We acknowledge Martin Eilers for feedback and Cato Stoffer for technical support. Funding was provided by the German Cancer Aid, Mildred-Scheel Early Career Center for Cancer Research, grant 8606100-NG1 (K.B.), the German Research Foundation, grant BA 7941/1-1 (A.B.), the European Research Council, grant TarMyc (E.W.) and the Helmholtz Young Investigator Group programme (M.M.). This publication was supported by the Open Access Publication Fund of the University of Würzburg.

## Author contributions

B.T. and K.B. conceived the project and performed the bulk of experiments. V.M. performed immunoblotting and imaging. G.C., J.H., P.B., C.P.A. and S.G. supported library preparations; G.C., A.B., Y.W., C.P.A., D.S. and P.G. performed bioinformatic analysis. E.W., M.M., and K.B. supervised the project. B.T. and K.B. wrote the draft. K.B. finalised the manuscript.

