## Supplementary material for "Nucleolar detention of NONO shields DNA double-strand breaks from aberrant transcripts": Supplemental_Fig_S1.pdf

### Trifault_Supplemental Fig. S1

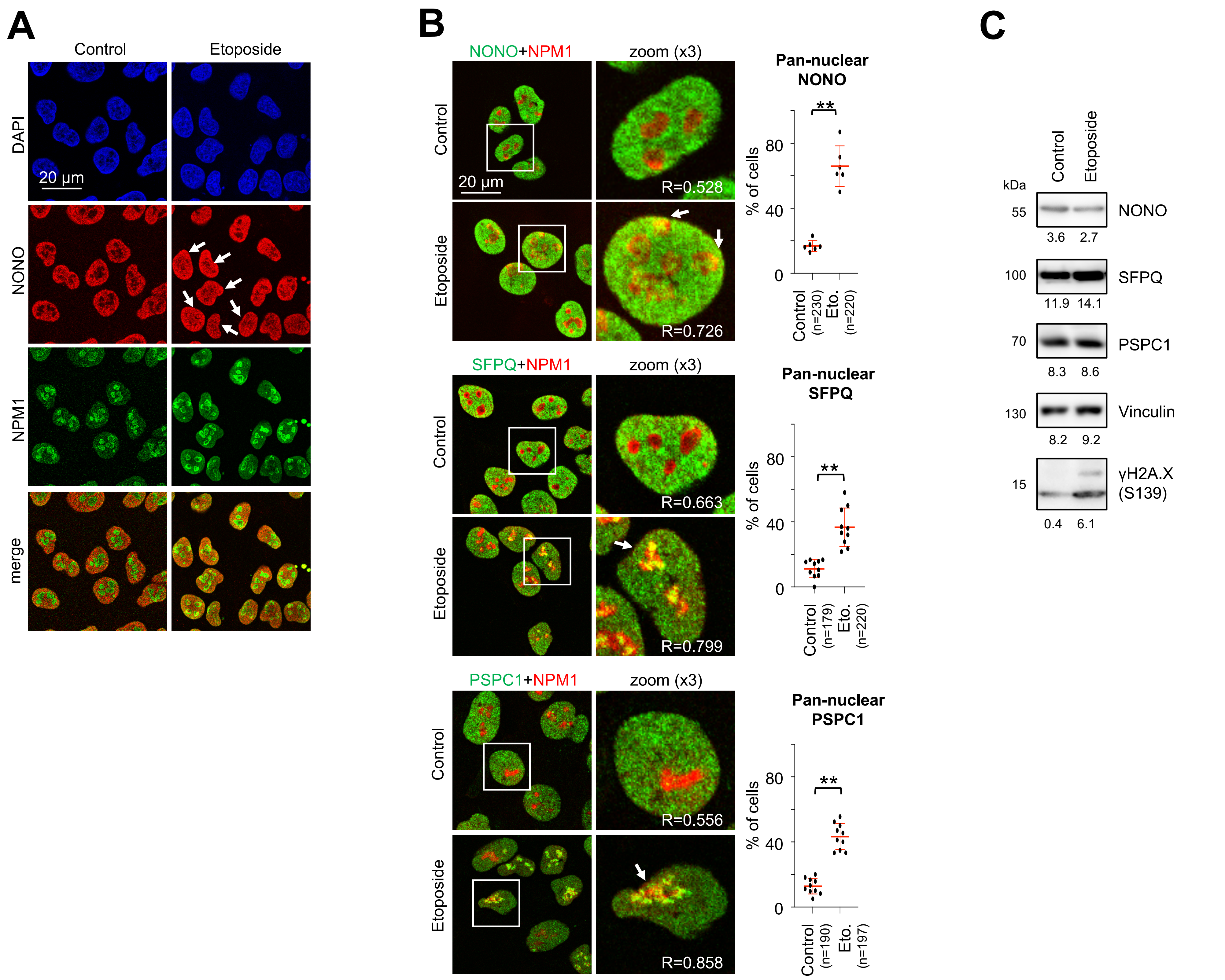


**Supplemental Figure S1.** Validation of NONO nucleolar re-localisation upon DNA damage in U2OS cells. (*A*) Imaging of non-POU domain containing octamer-binding protein (NONO) and nucleophosmin (NPM1) ±etoposide. Arrowhead, pan-nuclear NONO localisation. (*B*) Imaging (left) and quantitation (right) of NONO, splicing factor proline and glutamine rich (SFPQ), and paraspeckle component 1 (PSPC1) with NPM1 in the absence or presence (±) of etoposide. White box, zoom; arrowhead, colocalisation; R=Pearson correlation; n, number of cells analysed. Each dot represents % of cells with pan-nuclear signals as average from one acquisition. Arrowhead, pan-nuclear localisation. (*C*) Immunoblots detecting NONO, SFPQ, PSPC1 and ser-139 phosphorylated histone H2A.X variant (γH2A.X). Vinculin, loading control. Representative images are shown.
