## Supplementary material for "Nucleolar detention of NONO shields DNA double-strand breaks from aberrant transcripts": Supplemental_Fig_S2.pdf

### Trifault_Supplemental Fig. S2

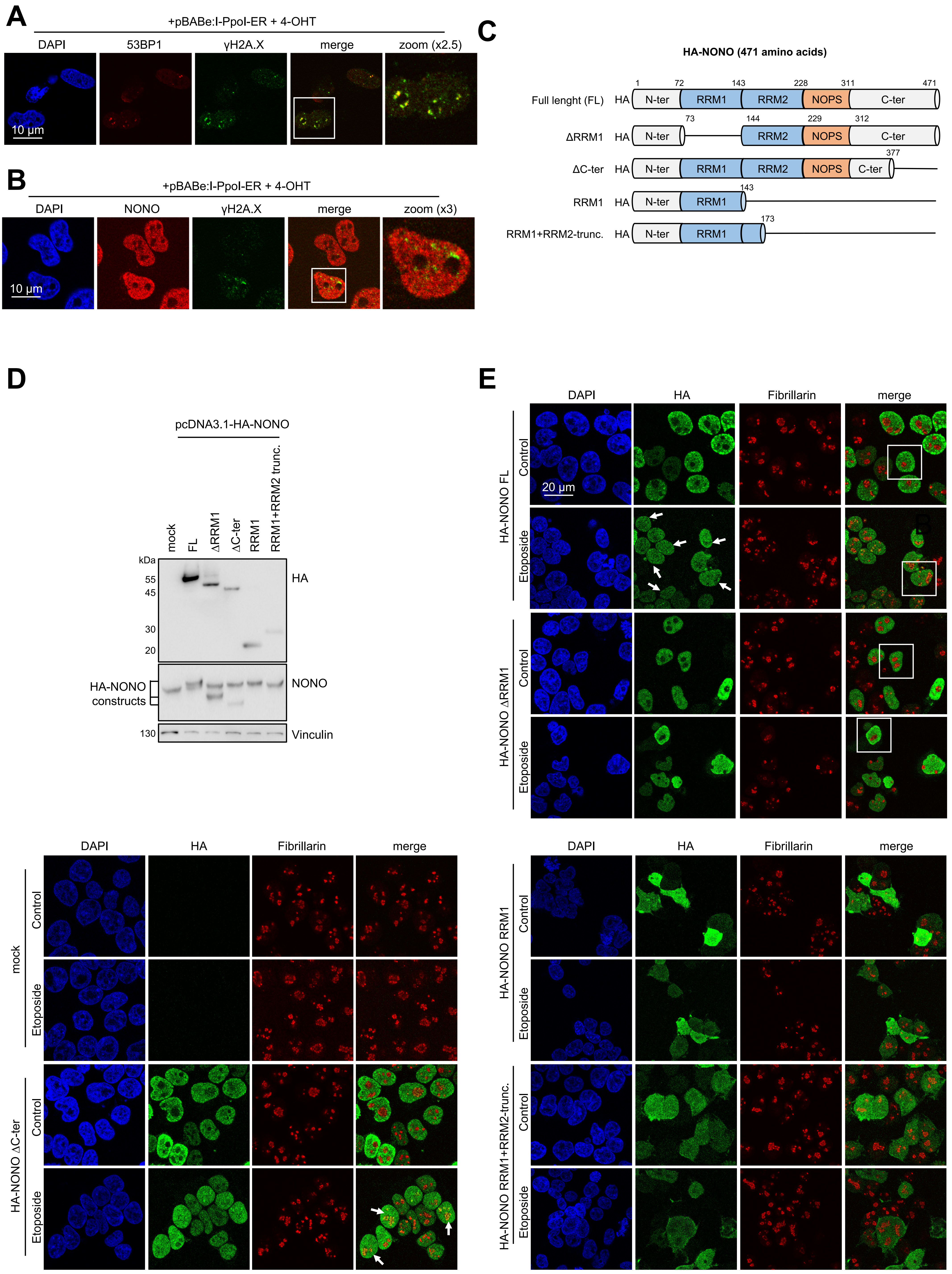

**Supplemental Figure S2.** Assessment of NONO localisation upon I-PpoI cleavage and HA-NONO constructs upon etoposide treatment in U2OS cells. (*A,B*) Imaging of 53BP1 and γH2A.X (*A*) or NONO and γH2A.X (*B*) upon transfection of I-PpoI encoding plasmid in U2OS cells in the presence of 4-hydroxytamoxifen (4-OHT). White box, zoom. (*C*) Scheme displaying the structure of HA-NONO expression constructs. RRM1, RNA recognition motif 1; RRM2, RNA recognition motif 2; NOPS, NonA/paraspeckle domain. (*D*) Immunoblots detecting HA-NONO variants and endogenous NONO. Vinculin, loading control; mock, non-transfected control. (*E*) Imaging of HA-NONO variants and fibrillarin ±etoposide. Mock, non-transfected control. Arrowhead, colocalisation. Arrowhead, pan-nuclear signal. Representative images are shown.
