## Supplementary material for "Nucleolar detention of NONO shields DNA double-strand breaks from aberrant transcripts": Supplemental_Fig_S3.pdf

### Trifault_Supplemental Fig. S3

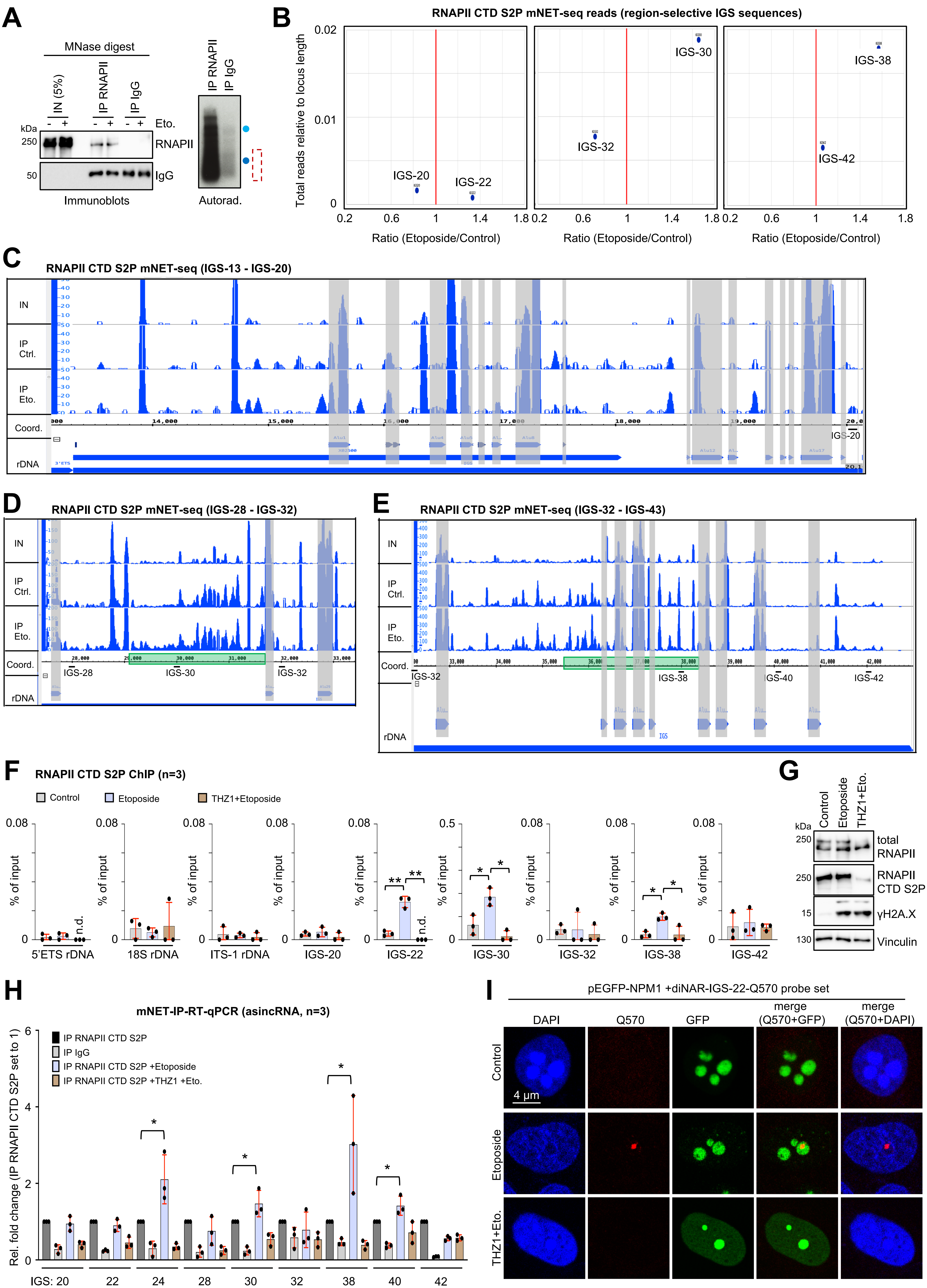

**Supplemental Figure S3.** Quality control for mNET-seq in U2OS cells. (*A*) Immunoblots detecting total RNAPII in input (IN, 5% of digest) and mNET-IP samples (10% material) that were immunoselected using RNAPII antibody from micrococcal nuclease (MNase)-digested samples ±etoposide; IgG, control (left). Autoradiograph detecting end-labeled transcripts upon immunoprecipitation (10% material) and PAGE separation (right). Blue dots, xylene cyanol/ bromophenol blue, size markers; red box, size-selected region. (*B*) Scatter plot displaying the number and relative abundance of mNET-seq reads ±etoposide (pairwise comparison of selected IGS sequences). (*C-E*) mNET-seq browser tracks for IGS consensus regions 13-20 (*C*), 28-32 (*D*), and 32-43 (*E*) from inputs (IN, merged) or after immunoprecipitation (IP) with CTD S2P-selective antibody ±etoposide. Red area, *Alu* element; green area, region of induction. IGS probe positions are not in scale. (*F*) CTD S2P ChIP with site-specific primers. n=number of biological replicates. (*G*) Immunoblots detecting total RNAPII, CTD S2P and γH2A.X. (*H*) RT-qPCR of transcripts associated with CTD S2P after immunoprecipitation (mNET-IP) ±etoposide or after preincubation with THZ1; IgG, control. (*I*) Imaging of GFP-NPM1 and RNA-FISH signals originating at IGS-22 using locus-specific, Quasar (Q)570-labeled RNA-FISH probes (n=30) ±etoposide or preincubation with THZ1. *, p-value <0.05; **, p-value <0.001; two-tailed t-test. Error bar, mean ±SD. Representative images are shown.
