## Supplementary material for "Nucleolar detention of NONO shields DNA double-strand breaks from aberrant transcripts": Supplemental_Fig_S4.pdf

### Trifault_Supplemental Fig. S4

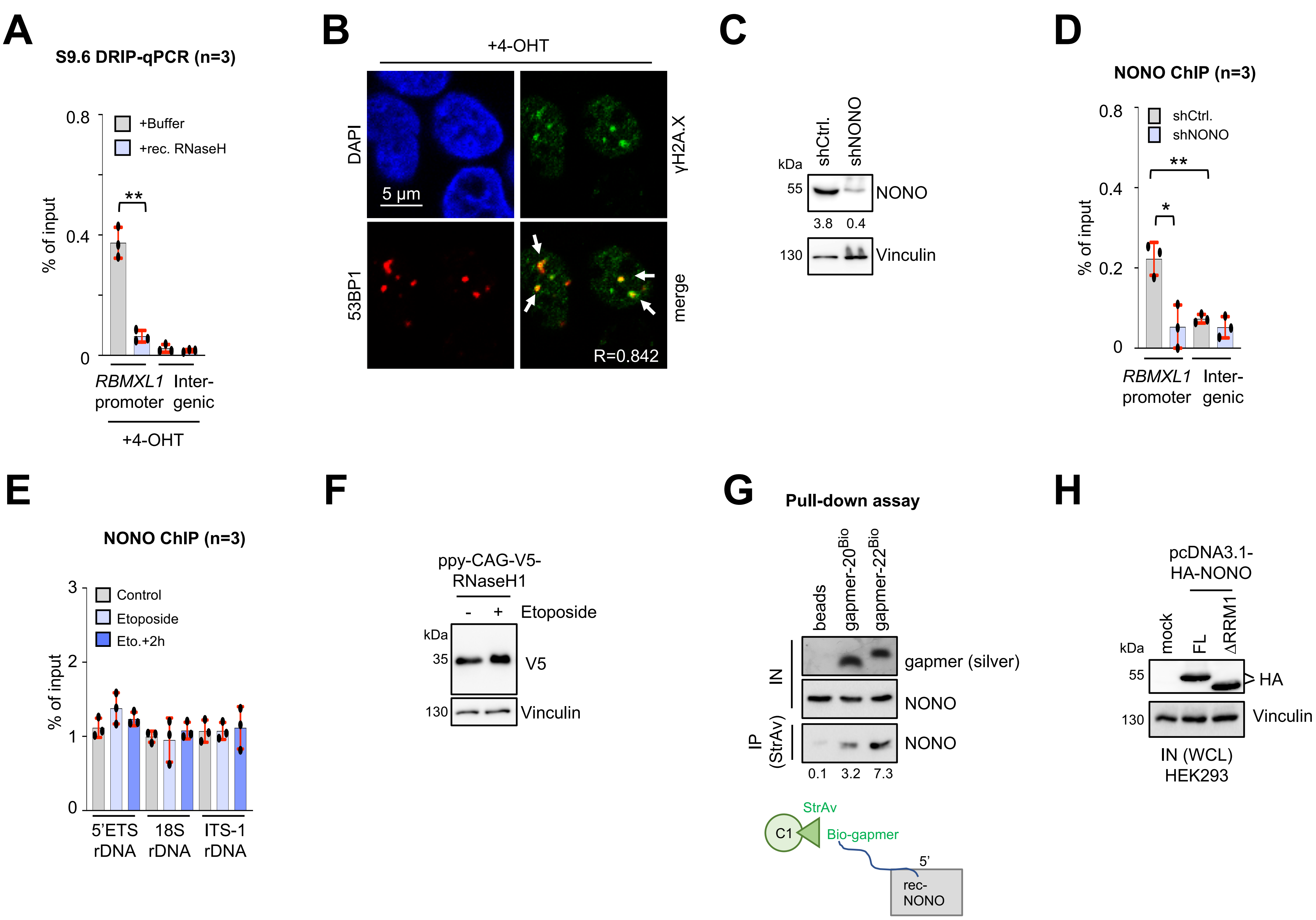


**Supplemental Figure S4.** Quality control for the assessment of R-loops, AsiSI restriction, NONO depletion and chromatin immunoprecipitation of NONO in U2OS cells as well as pull-down assay controls. (*A*) Quantitative PCR of DNA immunopurified from DNA-RNA hybrids (DRIP-qPCR) upon incubation with 4-OHT and recombinant RNaseH or buffer using S9.6 antibody and region-specific primers. (*B*) Imaging of 53BP1 and γH2A.X upon incubation with 4-OHT. Arrowhead, colocalisation. R=Pearson correlation. (*C*) Immunoblots detecting NONO upon shRNA transduction. Vinculin, loading control. (*D,E*) ChIP analysis of NONO occupancy using site-specific primers upon shRNA transduction (*D*) or ±etoposide or after chase (+2h) (*E*); 5'ETS, 5'external transcribed spacer; ITS-1, internal transcribed spacer-1. (*F*) Immunoblots detecting V5-RNaseH1 ±etoposide. Vinculin, loading control. (*G*) Pull-down assay displaying recombinant (rec)-NONO by immunoblotting after immunoprecipitation (IP) with biotin end-labeled (Bio) and immobilised gapmers. Silver stain and immunoblot of input (IN), loading controls, StrAv, streptavidin. (*H*) Immunoblots detecting HA-NONO variants FL and ΔRRM1 from HEK293 whole cell lysate (WCL) inputs (IN). Vinculin, loading control; mock, non-transfected control. *, p-value <0.05; **, p-value <0.001; two-tailed t-test. Error bar, mean ±SD. Representative images are shown.
