## Supplementary material for "Nucleolar detention of NONO shields DNA double-strand breaks from aberrant transcripts": Supplemental_Fig_S5.pdf

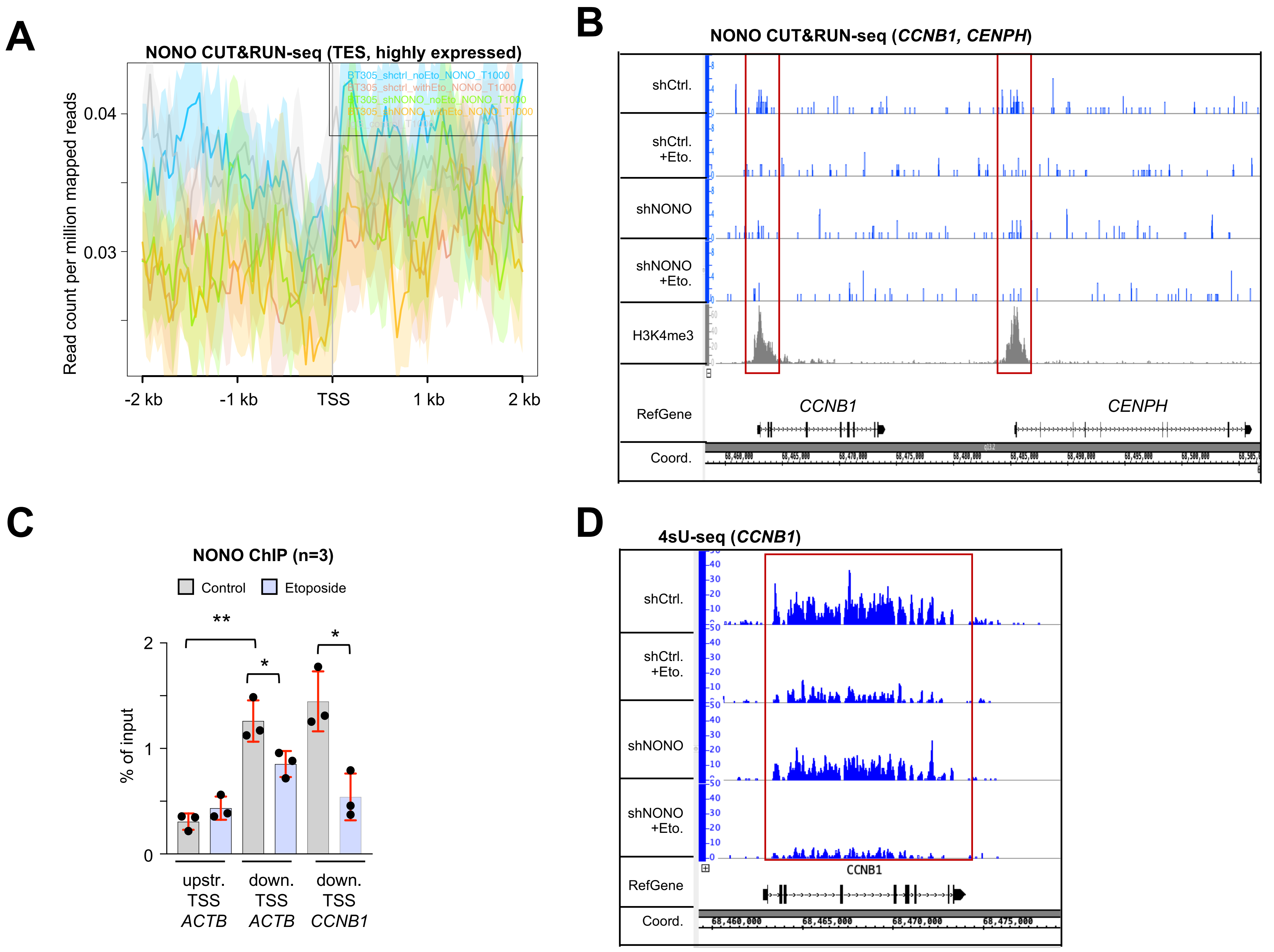


**Supplemental Figure S5.** DNA damage reduces NONO chromatin occupancy at TSSs of highly expressed genes and attenuates RNAPII activity in U2OS cells. (*A*) CUT&RUN-seq metagenes displaying NONO chromatin occupancy at transcriptional exit site (TES) of highly expressed genes ±NONO depletion/etoposide. (*B*) Browser tracks of NONO and histone H3 lys-4 tri-methylation (H3K4me3) CUT&RUN-seq ±NONO depletion/etoposide. Red box/H3K4me3, promoter region. (*C*) ChIP analysis of NONO occupancy using site-specific primers ±etoposide. (*D*) Browser tracks of 4sU-seq ±NONO depletion/etoposide. *, p-value <0.05; **, p-value <0.001; two-tailed t-test. Error bar, mean ±SD; red box, repressed region. Representative images are shown.
