## Supplementary material for "Nucleolar detention of NONO shields DNA double-strand breaks from aberrant transcripts": Supplemental_Fig_S6.pdf

### Trifault_Supplemental Fig. S6

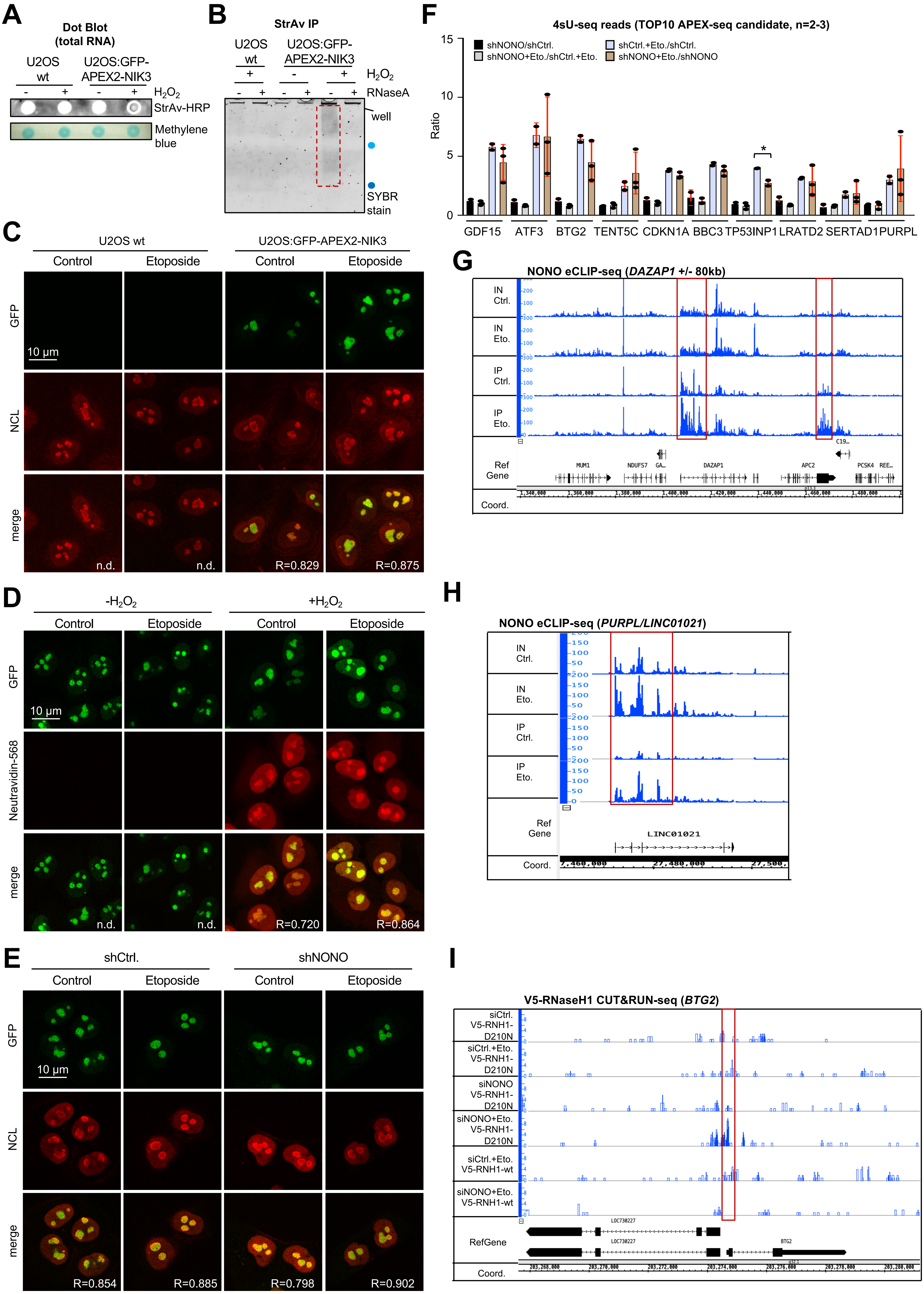

**Supplemental Figure S6.** Quality controls for APEX-seq, eCLIP-seq and detection of R-loops. (*A*) Dot blot analysis of total RNA from U2OS wild type (wt) or U2OS:GFP-APEX2-NIK3 cells using immunoblotting with a streptavidin-horseradish peroxidase (StrAv-HRP) probe ±hydrogen peroxide ($H_2O_2$). Methylene blue, loading control. (*B*) SYBR gold stain of RNA upon StrAv IP ±RNaseA digestion and PAGE separation from wt and U2OS:GFP-APEX2-NIK3 cells incubated ±$H_2O_2$. Blue dots, xylene cyanol/ bromophenol blue, size markers; red box, biotinylated material. (*C,D*) Imaging of GFP and NCL (*C*) in wild type and U2OS:GFP-APEX2-NIK3 cells ±etoposide or GFP and neutravidin-568 (*D*) in U2OS:GFP-APEX2-NIK3 cells ±etoposide and ±$H_2O_2$. (*E*) Imaging of GFP and NCL in U2OS:GFP-APEX2-NIK3 cells ±NONO depletion/etoposide. R=Pearson correlation; n.d., not detected; arrowhead, pan-nuclear NONO localisation. Representative images are shown. (*F*) Plot showing ratios of 4sU-seq reads for top 10 APEX-seq candidates ±NONO depletion/etoposide. (*G*) Browser tracks depicting NONO eCLIP-seq reads for the DAZAP1 transcript. Red box, increased binding. (*H*) Browser tracks depicting NONO eCLIP-seq reads for PURPL/LINC01021. Red box, increased binding. (*I*) Browser tracks depicting V5-RNaseH1 CUT&RUN-seq reads for *BTG2* ±NONO depletion/etoposide. Red box, region of increase.
