## Supplementary material for "Nucleolar detention of NONO shields DNA double-strand breaks from aberrant transcripts": Supplemental_Fig_S7.pdf

### Trifault_Supplemental Fig. S7

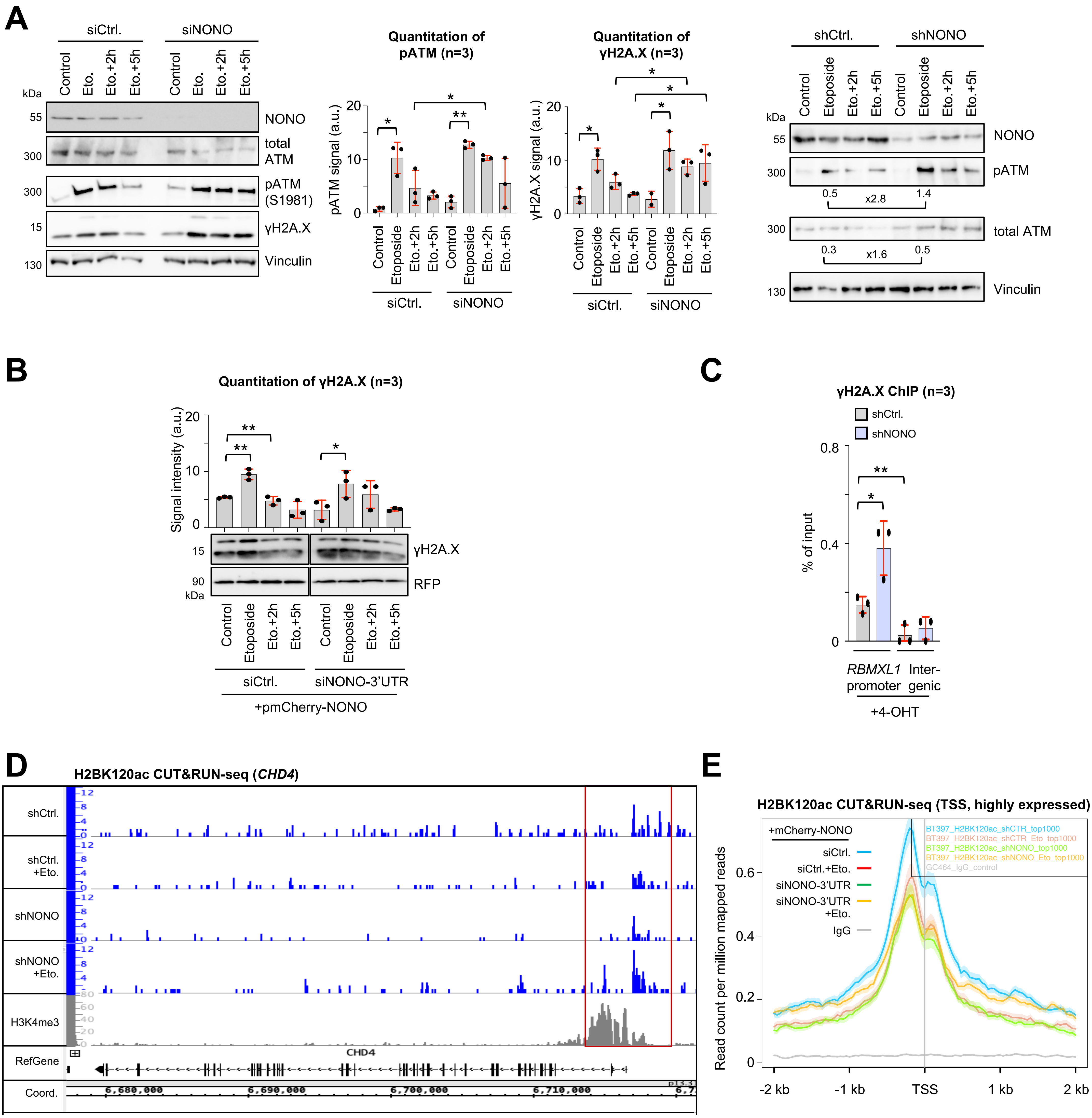


**Supplemental Figure S7.** Defects in DSB signaling upon depletion of NONO in U2OS cells. (*A*) Immunoblots (left) and quantitation (middle) of NONO, total ATM, phospho-(p)ATM and γH2A.X. ±NONO depletion/etoposide via siRNA (left) or shRNA (right); a.u., arbitrary units. Vinculin, loading control. (*B*) Quantitation (top) and immunoblots (bottom) detecting γH2A.X and RFP ±NONO depletion/etoposide and transient expression of mCherry-NONO. a.u., arbitrary units. (*C*) ChIP analysis of γH2A.X occupancy using site-specific primers upon 4-OHT incubation ±NONO depletion. (*D*) Browser tracks of histone H2B lys-120 acetylation (H2BK120ac) CUT&RUN-seq ±NONO depletion/etoposide for *CHD4*. Red box/H3K4me3, promoter region. (*E*) CUT&RUN-seq metagenes displaying histone H2B lys-120 acetylation (H2BK120ac) chromatin occupancy at TSSs of top 1000 highly expressed genes ±NONO depletion/etoposide and upon reexpression of mCherry-NONO. *, p-value <0.05; **, p-value <0.001; two-tailed t-test. Error bar, mean ±SD.
