## Supplementary material for "Nucleolar detention of NONO shields DNA double-strand breaks from aberrant transcripts": Supplemental_Table_S1.pdf

**Supplemental Table S1. Expression plasmids used in this study.**

| Plasmid | Source |
| --- | --- |
| pBABe:I-PpoI-ER | kind gift from Michael Kastan |
| pEGFP-NPM1 | kind gift from Xin Wang |
| pEGFP-RNaseH1 | kind gift from Martin Reijns |
| ppyCAG-V5-RNaseH1 wt | kind gift from Xiang-Dong Fu |
| ppyCAG-V5-RNaseH1 D210N | kind gift from Xiang-Dong Fu |
| pmCherry-NONO | kind gift from Ling-Ling Chen |
| pcDNA3.1-HA-NONO | kind gift from Nicolas Manel |
