## Supplementary material for "Nucleolar detention of NONO shields DNA double-strand breaks from aberrant transcripts": Supplemental_Table_S2.pdf

**Supplemental Table S2. Primer pairs used for site-directed mutagenesis and siRNA or shRNA for RNA interference.**

| **Primer** | **Sequence (5'-3')** |
|---|---|
| ΔRRM1-FWD | GCCTGCCATAGTGCATCCCTTAC |
| ΔRRM1-REV | TTGGGTGAAGGTCTTCTCTCCTGGTTTTC |
| ΔC-TER-FWD | TAACTCGAGCATGCATCTAGAGGGC |
| ΔC-TER-REV | CGCATCAGGGAAGGTTCCC |
| RRM1-FWD | TAACTCGAGCATGCATCTAGAGGGC |
| RRM1-REV | AAAGCGCACACGCAGCTGC |
| RRM1+RRM2-trunc-FWD | TAACTCGAGCATGCATCTAGAGGGC |
| RRM1+RRM2-trunc-REV | CTACAGCCCTCTCTACCTG |
| **siRNA** | **Sequence (5'-3')** |
| siCtrl. | smart pool |
| siNONO | smart pool |
| siNONO-3'UTR | GGAGUAUGCUGGAGGCAGAdTdT |
| **shRNA** | **Source** |
| pLKO.1-puro-NONO-sh1 | kind gift from Nicolas Manel |
| pLKO.1-puro-NONO-sh4 | kind gift from Nicolas Manel |
| pLKO.1-puro-non-target shRNA, shCtrl. | Sigma |
