## Supplementary material for "Nucleolar detention of NONO shields DNA double-strand breaks from aberrant transcripts": Supplemental_Table_S3.pdf

**Supplemental Table S3. Antibodies used in this study.**

| **Primary antibody** | **Species** | **Supplier, code** |
|---|---|---|
| IgG control | rabbit | Proteintech, 30000-0-AP |
| Anti-NONO | rabbit | Proteintech, 11058-1-AP |
| Anti-PSPC1 | rabbit | Proteintech, 16714-1-AP |
| Anti-SFPQ [EPR11874] | rabbit | Abcam, ab177149 |
| Anti-NPM1 [FC82291] | mouse | Abcam, ab10530 |
| Anti-phospho-histone H2A.X (S139) | rabbit | Cell Signaling, 2577 |
| Anti-phospho-histone H2A.X (S139) [JBW301] | mouse | Millipore, 05-636 |
| Anti-nucleolin [364-05] | mouse | Abcam, ab136649 |
| Anti-fibrillarin | rabbit | Abcam, ab5821 |
| Anti-vinculin | mouse | Sigma, V9131 |
| Anti-histone H2B | rabbit | Abcam, ab1790 |
| Anti-HA tag [16B12] | mouse | BioLegend, 901502 |
| Anti-HA tag | rabbit | Abcam, ab9110 |
| Anti-RNA polymerase II RPB1 | mouse | BioLegend, 920204 |
| Anti-RNA polymerase II CTD repeat YSPTSPS | rabbit | Abcam, ab26721 |
| Anti-phospho-RNA polymerase II CTD repeat YSPTSPS (S2) | rabbit | Abcam, ab5095 |
| Anti-V5 tag | mouse | Thermo, R960-25 |
| Anti-SPT5 [D-3] | mouse | Santa Cruz, sc133217 |
| Anti-DNA-RNA-hybrid [S9.6] | mouse | Millipore, MABE1095 |

| | | |
|---|---|---|
| Anti-53BP1 | rabbit | Novus, NB100-304 |
| Anti-histone H3K4me3 | rabbit | Abcam, ab8580 |
| Anti-phospho-ATM (S1981) [EP1890Y] | rabbit | Abcam, ab81292 |
| Anti-phospho-ATM/ATR substrate (S*Q) [D23H2/D69H5] | rabbit | Cell Signaling, 9607 |
| Anti-histone H2BK120ac | rabbit | Active Motif, 39120 |
| Anti-RFP tag [RF5R] | mouse | Thermo, MA5-15257 |
| Anti-GFP tag | rabbit | Abcam, ab290 |
| Anti-ATM [D2E2] | rabbit | Cell Signaling, 2873 |
| **Secondary antibody** | **Species** | **Supplier, code** |
| HRP-linked IgG | mouse | Cytivia, NA931 |
| HRP-linked IgG | rabbit | Cytivia, GEHENA934 |
| Alexa Fluor 546-linked IgG | rabbit | Thermo, A10040 |
| Alexa Fluor 488-linked IgG | mouse | Thermo, A32766TR |
| Alexa Fluor 546-linked IgG | mouse | Thermo, A10036 |
| Alexa Fluor 488-linked IgG | rabbit | Thermo, A21206 |
