## Supplementary material for "Nucleolar detention of NONO shields DNA double-strand breaks from aberrant transcripts": Supplemental_Table_S4.pdf

**Supplemental Table S4. Gapmers used for end-labeling and binding assays.** All bases are DNA except ones preceded by 'm' like mU, which are 2'-hydroxy methylated RNA bases. *, phosphorothioate linkage. Gapmers were custom made (IDT).

| **Gapmer** | **sequence (5'-3')** |
|---|---|
| gapmer-20 | mG*mU*mA*mG*mC*C*T*T*G*G*G*C*T*T*C*mU*mC*mU*mC*mC |
| gapmer-22 | mC*mA*mG*mU*mG*G*C*T*C*A*C*G*T*C*T*mG*mU*mC*mA*mU |
