## Supplementary material for "Nucleolar detention of NONO shields DNA double-strand breaks from aberrant transcripts": Supplemental_Table_S5.pdf

**Supplemental Table S5. Adapters used for DSB ligation.**

| BLISS adapter | Sequence (5'-3') |
|---|---|
| adapter A1 | CATCACGC |
| adapter A2 | GTCGTTCC |
| adapter A3 | TGATGATC |
| adapter A4 | ACGACATC |
