## Supplementary material for "Nucleolar detention of NONO shields DNA double-strand breaks from aberrant transcripts": Supplemental_Table_S6.pdf

**Supplemental Table S6. Primer pairs used for RT-qPCR and ChIP.**

| **Primer** | **Sequence (5'-3')** |
|---|---|
| IGS20-fwd | GTAGCCTTGGGCTTCTCTCC |
| IGS20-rev | AGTTTTCAGCCCCAACACAC |
| IGS22-fwd | CAGTGGCTCACGTCTGTCAT |
| IGS22-rev | CGCCTGACTCCATTTCGTAT |
| IGS24-fwd | CCCGCGCACATAATAACTAA |
| IGS24-rev | AAATCACTCCTCACGGGAAC |
| IGS28-fwd | CCTTCCACGAGAGTGAGAAG |
| IGS28-rev | GACCTCCCGAAATCGTACAC |
| IGS30-fwd | GGTCTCTGCGTCTCGCTATC |
| IGS30-rev | TGAAGAATTCAGGCCTTGGT |
| IGS32-fwd | AAAAGCTGGCCGATCTGAAT |
| IGS32-rev | CGTCTGTTCAGCTATTTTGCAG |
| IGS38-fwd | CTCACAGAGGAAGGGAGCAC |
| IGS38-rev | AACAAGGGAGGGAGGAACTT |
| IGS40-fwd | TTCTCCTTGGTCAGGGGTTT |
| IGS40-rev | CAGGAAAGTCCCCAACAACA |
| IGS42-fwd | GCTTCTCGACTCACGGTTTC |
| IGS42-rev | CCGAGAGCACGATCTCAAAG |
| 5‘ETS-fwd | GCCCCGGGGGAGGTAT |
| 5‘ETS-rev | GAGGACAGCGTGTCAGC |
| 18S-fwd | GTTGAACCCCATTCGTGATG |
| 18S-rev | GGGACTTAATCAACGCAAGC |

| | |
|---|---|
| ITS-1-fwd | TGTGAAACCTTCCGACCCC |
| ITS-1-rev | GGGGTTGCCTCAGGCC |
| upstream-TSS-ACTB-fwd | CCCACCTGACAACCTCTCAT |
| upstream-TSS-ACTB-rev | CCCTTCTTGCTGCCTGTT |
| downstream-TSS-ACTB-fwd | CTCAATCTCGCTCTCGCTCT |
| downstream-TSS-ACTB-rev | CTCGAGCCATAAAAGGCAAC |
| downstream-TSS-CCNB1-fwd | ATCGCCCTGGAAACGCATTCT |
| downstream-TSS-CCNB1-rev | GCCAGCCTAGCCTCAGATTTA |
| RBMXL1-promoter-DSB-fwd | GATTGGCTATGGGTGTGGAC |
| RBMXL1-promoter-DSB-rev | CATCCTTGCAAACCAGTCCT |
| Intergenic-no DSB-fwd | ATTGGGTATCTGCGTCTAGTGAGG |
| Intergenic-no DSB-rev | GACTCAATTACATCCCTGCAGCT |
