## Supplementary material for "Nucleolar detention of NONO shields DNA double-strand breaks from aberrant transcripts": Supplemental_Table_S7.pdf

**Supplemental Table S7. Quasar-570-labeled probes used for RNA-FISH.**

| Probe | Sequence (5'-3') |
|---|---|
| IGS-22 (30 probes) | aaatgaaagaaaaggcac |
| | acagatatacaagaaaga |
| | catacatacatccatgca |
| | catacatacatacataca |
| | tccagataaatacgtaca |
| | gacagagtgagacccggt |
| | actgcactacagcctggg |
| | agtgacccaagatcgcac |
| | cctggcaggcggaggctg |
| | ggtggaagaattgcttga |
| | cagctactcgggaggctg |
| | ggggcaggcacctgtaac |
| | aaatggagtcaggcgccg |
| | cgtctctactgaaaatac |
| | ggccaacgtggtgaaacc |
| | aggagttcgagaccagcc |
| | cgtctgtcatcccgaggt |
| | cactgaacgcagtggctc |
| | tctaagaaatggtactgt |
| | tgttcccgtgagagtgat |
| | cacgtcacccataagtgt |
| | taactaactaaaatctct |
| | ccacctcctcgcgcacat |

| | |
|---|---|
| | agcaagagccaaactccg<br>cattgcactgtagcctgg<br>actccttgagtcccctag<br>gtgtcctctctgccgtag<br>cccttacgctcagaatga<br>ctgccacctttcgctgtg<br>agcctttaaaagcgcggc |
